# How good a predictor the *in vitro* antibiogram is, of *in vivo* antibiotic efficacy? An *in vitro* - *in vivo* integrated approach

**DOI:** 10.64898/2026.11.17.766954

**Authors:** Mitva Vaghela, Nidhi Thakkar, Bruce Blough, Vijay Kothari

## Abstract

Conventional antimicrobial susceptibility testing, particularly the disc diffusion assay (DDA), provides a convenient means of assessing bacterial susceptibility to antibiotics. However, its ability to predict antibiotic efficacy in an infected host remains uncertain. This study investigated the relationship between DDA-derived inhibition zones and host-protective antibiotic activity using *Caenorhabditis elegans* infection models involving three bacterial pathogens: methicillin-resistant *Staphylococcus aureus* (MRSA), *Pseudomonas aeruginosa*, and *Clostridioides difficile*. A total of 42 antibiotics were evaluated through DDA, while a separate panel of 75 antibiotics was screened using an automated *C. elegans* infection assay. Twenty-six antibiotics common to both panels were selected for comparative analysis. Gnotobiotic L3-L4 stage worms were challenged with individual pathogens in the presence of each antibiotic at a fixed concentration of 9.6 µg/mL. Worm activity was quantified at 24, 48, and 72 h using an automated worm tracker system, with appropriate health, infection, vehicle, and antibiotic-toxicity controls.

Overall, DDA-derived zone of inhibition (ZOI) diameters showed weak-to-moderate, pathogen- and time-dependent correlations with worm activity. At 24 h, Pearson’s correlation coefficients ranged from −0.20 to 0.31 across the three pathogen-host systems, while at 48 h they ranged from −0.009 to 0.49. Notably, antibiotic rankings based on DDA and host-protective activity differed considerably. For example, against *C. difficile*, imipenem produced the largest inhibition zone (41 mm) but failed to protect worms at 24 and 48 h, whereas tobramycin, despite producing a smaller inhibition zone (26 mm), exhibited the highest early worm activity. Similar discrepancies were observed with the other pathogens.

These findings demonstrate that DDA-derived susceptibility profiles do not consistently predict antibiotic-mediated protection in an infected host. The *C. elegans* model may therefore serve as a complementary whole-organism screening platform, providing additional functional information for prioritizing antimicrobial agents beyond conventional *in vitro* susceptibility testing.

## 1. Introduction

The rapid emergence of antimicrobial resistance among bacterial pathogens necessitates highly accurate screening methods to guide effective clinical interventions. Currently, the antibiogram generated through the disc diffusion assay (DDA) is widely used to assess the antibiotic susceptibility of clinical isolates of pathogenic bacteria. This standard *in vitro* methodology provides rapid, cost-effective profiling by measuring the diameter of the zone of inhibition to classify bacterial isolates as susceptible, intermediate, or resistant. However, there is a vast difference between the matrix environment of this kind of *in vitro* assay and the complex physiological milieu present *in vivo*. Because of these distinct environmental and biological disparities, traditional standard susceptibility testing may not always correlate reliably with *in vivo* therapeutic success (Fantin et al., 1991; Blaser et al., 1995).

When *in vitro* susceptibility profiles fail to accurately mirror *in vivo* outcomes, it can lead to critical mismatches in therapeutic application. Agar-based assays like the DDA do not account for host-pathogen interactions, tissue penetration, or the immunomodulatory effects of the host environment (Blaser et al., 1995). Therefore, exploring alternative whole-organism platforms is essential to bridge the gap between initial screening and functional clinical verification (Kong et al., 2016). We propose that antibiotic susceptibility testing in a *Caenorhabditis elegans* platform is worthy of further investigation as a potentially more robust predictor of antibiotic efficacy for therapeutic purposes.

*C. elegans* has emerged as a highly versatile, scalable *in vivo* model host for studying bacterial pathogenesis, innate immunity, and host resilience (Zarroug et al., 2023). The nematode model bypasses certain ethical and financial constraints of mammalian lines while allowing for the continuous evaluation of physiological metrics during an active infection (Kong et al., 2016). In this study, our primary objective was to investigate the correlation between the antibiotic susceptibility observed in a DDA and the *in vivo* antibiotic efficacy observed in a worm model of infection. To ensure the biological relevance of our findings, we evaluated the *in vitro* antibiotic susceptibility profiles of three prominent pathogens-*Pseudomonas aeruginosa, Clostridioides difficile*, and methicillin-resistant *Staphylococcus aureus* (MRSA)- all of which have been studied in *C. elegans* infection and survival assays (Tan et al., 1999; Kim et al., 2020; Abouelkhair et al., 2026).

To rigorously quantify these outcomes, we challenged the host worms (25 worms of the L3-L4 stage per well) with either of these pathogens in the presence or absence of antibiotics. Rather than relying on simple endpoint lethality, worm activity was quantified over a three-day period through an automated worm tracker, serving as a comprehensive surrogate for their overall survival and healthspan. By analysing the correlation coefficient between the DDA zone of inhibition diameters and these *in vivo* efficacy metrics, this study aims to systematically evaluate the predictive reliability of standard *in vitro* antibiograms. Ultimately, highlighting the mismatches between agar-derived data and whole-organism survival will underscore the necessity of integrating complex *in vivo* models like *C. elegans* into the antimicrobial drug discovery and validation pipeline (Kong et al., 2016).

## 2. Methods

### 2.1 Antibiotics

For disc diffusion assay, the antibiotic disc panels (Icosa G-I PLUS; Icosa G-I MINUS; Dodeca Universal – I; Dodeca G – XI - MINUS) were procured from HiMedia, Mumbai, India. For *in vivo* assay, the panel of 75 antibiotics (in aluminium foil covered 96 well plate; RTI-CARB13-1-D31) was provided by RTI International under Chemistry Center for Combating Antibiotic Resistant Bacteria (CC4CARB) program. Each antibiotic was provided dissolved in DMSO at the concentration of 6.4 mg/mL. An aliquot of 1.5 µL from the antibiotic well was transferred into the 24-well plates (1 mL assay volume per well) used for *in vivo* assays, making the final antibiotic concentration in each well 9.6 ppm. Total number of non-overlapping and common antibiotics between discs and those used in *in vivo* assay was 91 and 26 respectively.

### 2.2 Test organisms

#### 2.2.1 Bacterial pathogens

The strain of *S. aureus* (ATCC 43300) utilized in this study was procured from HiMedia. This strain is claimed to be resistant to methicillin and oxacillin [https://www.atcc.org/products/43300; Accessed on 1 March 2024], and our transcriptome of this strain, conducted as a part of another study (Thakkar et al., 2024), showed this strain to be positive for expression of the *mecA* gene, whose presence is considered a defining characteristic for labelling *S. aureus* as methicillin-resistant (Martinez and Simonsen, 2017). Antibiogram of this strain generated in our lab showed it to be resistant to four macrolide antibiotics (erythromycin, clindamycin, azithromycin, and clarithromycin), and intermediate to amoxicillin/clavulanic acid.

The *P. aeruginosa* strain used in this study was sourced from our internal culture collection. This strain has been well characterized by us (Gajera et al., 2023) with respect to its antibiotic resistance, pigment production and certain other virulence traits. Its antibiogram generated through a disc-diffusion assay performed as per Clinical and Laboratory Standards Institute (CLSI) guidelines revealed apparent resistance to eight antibiotics: co-trimoxazole, Augmentin, nitrofurantoin, ampicillin, chloramphenicol, clindamycin, cefixime, and vancomycin, representing six broad pharmacological groups: sulfonamide combinations, β-lactams, nitrofurans, amphenicols, lincosamides, and glycopeptides. Hence, it can be described as a multidrug resistant (MDR) strain. As reported in our earlier publications (Gajera et al., 2023) involving this strain, it is a haemolytic strain capable of producing the quorum sensing (QS) regulated pigments (pyocyanin and pyoverdine), and also of biofilm formation.

The strain of *C. difficile* utilized in this study (ATCC 9689) was procured from HiMedia. This bacterium was cultured under anaerobic conditions in an anaerobic jar using an AnaeroGas Pack (HiMedia) with anaerobiosis confirmed using an Anaero Indicator tablet (HiMedia). In the disc diffusion assay, five antibiotics (cefpodoxime, cefixime, methicillin, oxacillin and colistin) produced no detectable zone of inhibition against *C. difficile*. However, since standardized disc-diffusion interpretive breakpoints are not established for this organism, the observed inhibition zones are presented as descriptive measurements rather than definitive susceptible/intermediate/resistant classifications (Table S1). This bacterium was maintained on modified reinforced clostridial agar (HiMedia) slants under anaerobic condition. For the purpose of different assays described in this report, (unless described otherwise) it was grown in modified reinforced clostridial broth (pH 7.0 ± 0.2) for 21 ± 1 h at 35 ± 0.5°C.

#### 2.2.2 Model host

Wild type N2 Bristol strain of *C. elegans* procured from the Caenorhabditis Genetics Center (Minneapolis, USA) was used as a model organism in this study. Lyophilized *E. coli* OP50 procured from Biovirid (Netherlands) was used as food for *C. elegans* while maintaining the worm on NGM agar plates (Nematode Growing Medium). Worm synchronization was done as described in literature (Stiernagle, 2006). Prior to the *in vivo* assays, worms were kept without food for two days to make them gnotobiotic.

### 2.3 Disc diffusion assay

Antibiogram of bacterial strains was obtained through a DDA performed in accordance with the CLSI guidelines (Lewis, 2024), where available. Each bacterial pathogen was grown on its specific agar media, and after overnight incubation 2-3 isolated colonies were used to prepare the inoculum for the subsequent disc diffusion assay by resuspending the colonies in normal saline. The OD_625_ of the inoculum was adjusted to range from 0.08–0.10 to achieve equivalence to McFarland turbidity standard 0.5. The known volume (0.1 mL) of this inoculum was then spread onto cation-adjusted Muller–Hinton agar (HiMedia) in 150 mm plates (Borosil, Mumbai, India), followed by loading antibiotic discs onto the agar surface. The plates were then incubated at 35 ± 0.5 °C for 18 ± 1 h, followed by the observation and measurement of the diameter of the zone of inhibition.

#### 2.4 *In vivo* antibiotic efficacy assay

Gnotobiotic *Caenorhabditis elegans* worms at the L3–L4 larval stage was collected from NGM agar plates and transferred into 24-well plates, with each well containing 25 ± 5 worms in a final assay volume of 1 mL. The worms were challenged with the respective bacterial pathogen in the presence or absence of the test antibiotic. Appropriate quantities of bacterial culture (100 µL; OD_764_=1.50 for *P. aeruginosa* and MRSA; OD_625_=1.50 for *C. difficile*), antibiotic stock solution and M9 buffer were added to achieve the desired experimental conditions, with a final antibiotic concentration of 9.6 µg/mL in each antibiotic-containing well. Bacterial culture to be used in this assay was obtained by growing (at 35 ± 0.5^0^C for 21 ± 1 h) MRSA, *P. aeruginosa*, and *C. difficile* in tryptone soy broth, pseudomonas broth (supplemented with 3%v/v glycerol), and modified reinforced clostridial broth respectively. For assays involving *Clostridioides difficile*, the perimeter of the 24-well plates was wrapped with Parafilm before incubation.

Four types of controls were included in the assay: (i) Health Control, consisting of worms exposed neither to the bacterial pathogen nor to the antibiotic; (ii) Infection Control, consisting of worms challenged with the bacterial pathogen in the absence of antibiotic; (iii) Vehicle Control, consisting of worms challenged with the bacterial pathogen in the presence of DMSO (0.15%, v/v); and (iv) Toxicity Control, consisting of worms exposed to the antibiotic in the absence of the bacterial pathogen. Each pathogen was capable of killing 100% of challenged worms within 24 h.

Worm Activity was monitored at 24, 48 and 72 h using an automated worm-tracking system (WMicrotracker® ARENA, Phylumtech, Argentina). The resulting activity counts were used as a quantitative measure of the motility-associated functional status of the worm population. Qualitative observations of worm survival, activity, and fertility were also recorded using a binocular microscope (LABOMED VISION 2000; 4× objective). Representative videos were captured using a Magnus camera (5.1 MP) attached to the microscope.

### 2.5 Statistics

All values reported are means of two or more independent assays, each of which contained two or three replicates. Comparison of worm activity between Antibiotic Well and Vehicle Control well was achieved through a two-tailed t-test, whereas efficacy of multiple antibiotics within a class was analysed through Tukey’s multiple comparisons test following two-way ANOVA (P < 0.05). Pearson’s correlation coefficient (r) shown in Table 1-3 was calculated using Microsoft Excel (version 2016) to assess the linear association between antibiotic inhibition-zone diameter and worm activity at each observation time point.

**Table 1.**
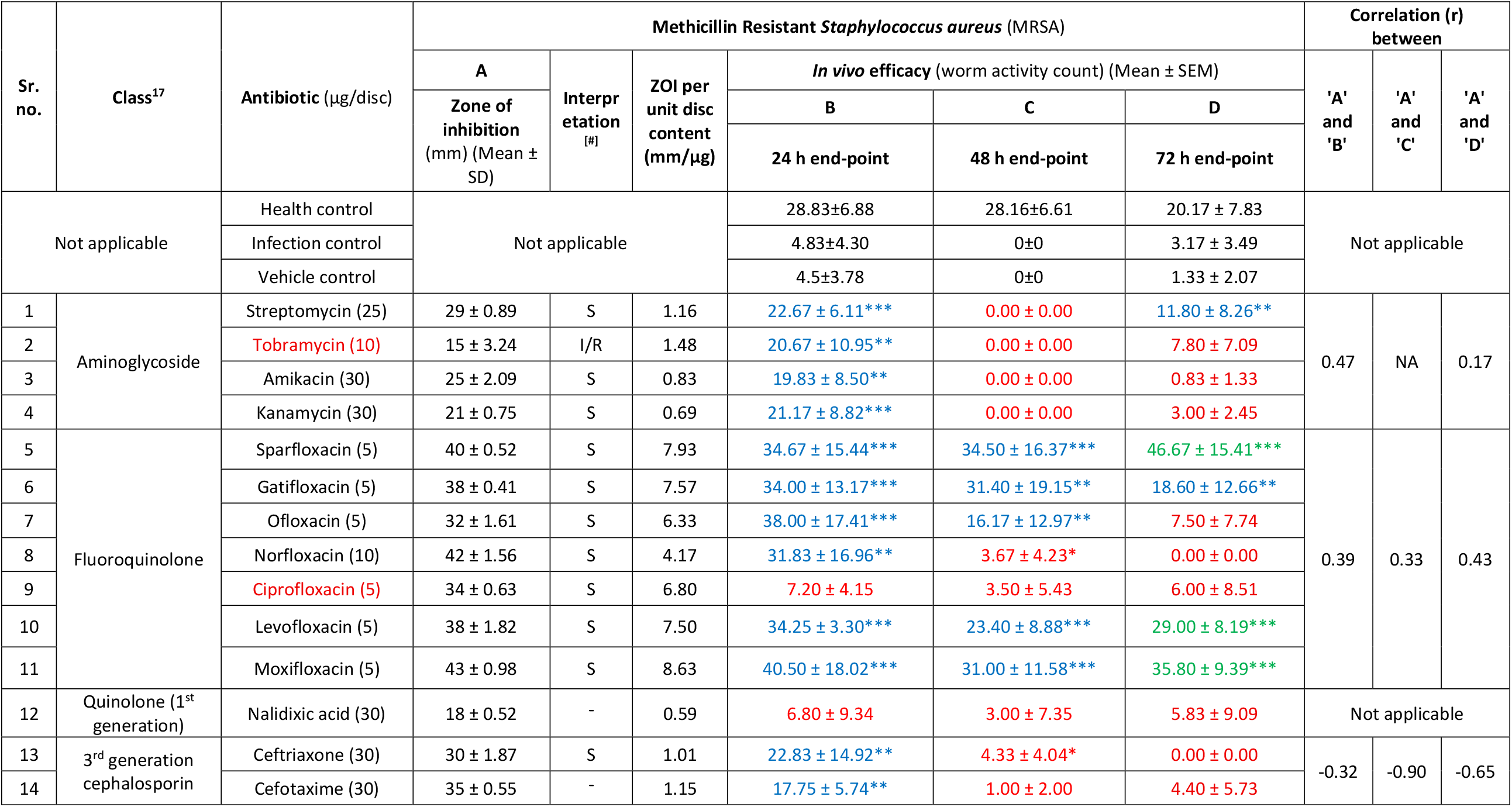
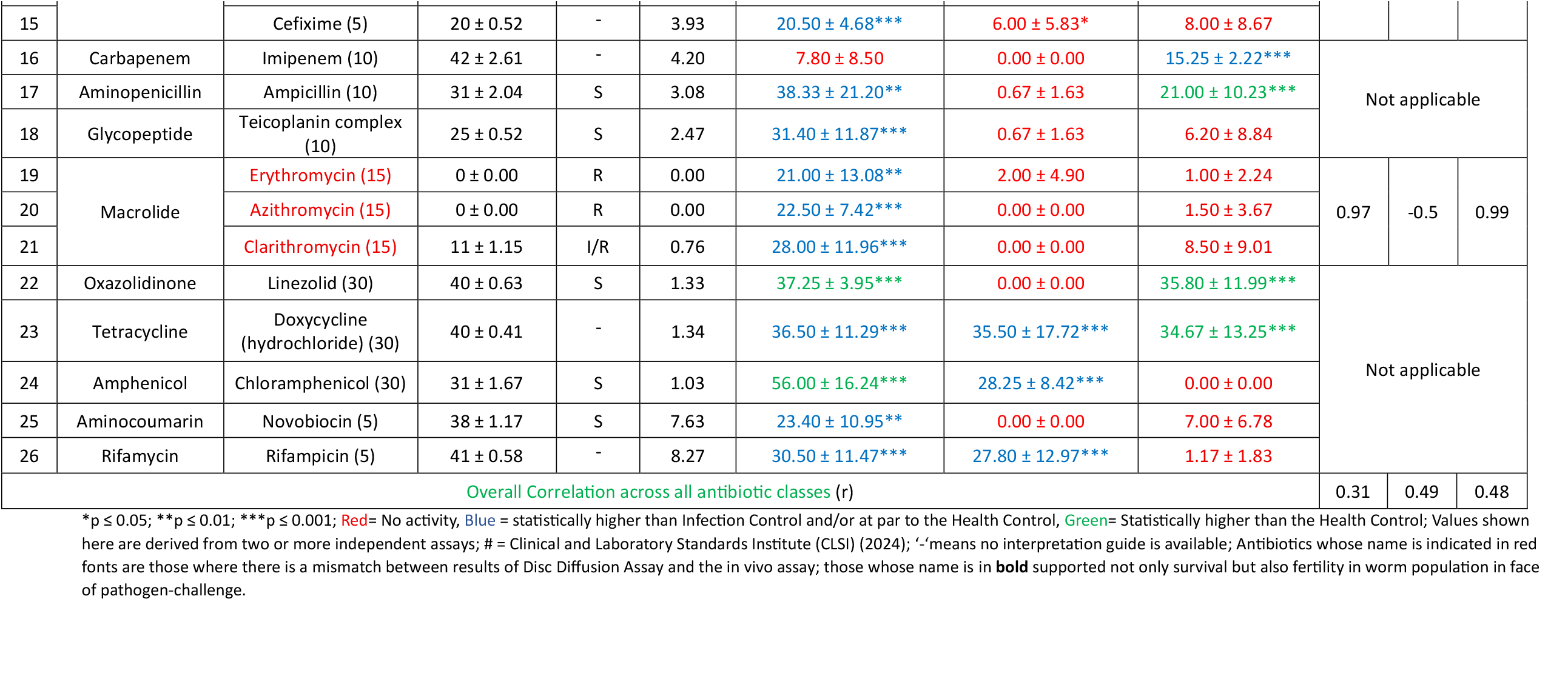
Comparison of *in vitro* and *in vivo* efficacy of antibiotics against MRSA.

| **Sr. no.** | **Class<sup>17</sup>** | **Antibiotic** (µg/disc) | **Methicillin Resistant *Staphylococcus aureus*** (MRSA) |  |  |  |  |  | **Correlation (r) between** |  |  |
|---|---|---|---|---|---|---|---|---|---|---|---|
|  |  |  | **A** | **Interpretation<sup>[#]</sup>** | **ZOI per unit disc content (mm/µg)** | ***In vivo* efficacy** (worm activity count) (Mean ± SEM) |  |  | **'A' and 'B'** | **'A' and 'C'** | **'A' and 'D'** |
|  |  |  | **Zone of inhibition** (mm) (Mean ± SD) |  |  | **B** | **C** | **D** |  |  |  |
|  |  |  |  |  |  | **24 h end-point** | **48 h end-point** | **72 h end-point** |  |  |  |
| Not applicable |  | Health control | Not applicable |  |  | 28.83±6.88 | 28.16±6.61 | 20.17 ± 7.83 | Not applicable |  |  |
|  |  | Infection control |  |  |  | 4.83±4.30 | 0±0 | 3.17 ± 3.49 |  |  |  |
|  |  | Vehicle control |  |  |  | 4.5±3.78 | 0±0 | 1.33 ± 2.07 |  |  |  |
| 1 | Aminoglycoside | Streptomycin (25) | 29 ± 0.89 | S | 1.16 | 22.67 ± 6.11*** | 0.00 ± 0.00 | 11.80 ± 8.26** | 0.47 | NA | 0.17 |
| 2 |  | Tobramycin (10) | 15 ± 3.24 | I/R | 1.48 | 20.67 ± 10.95** | 0.00 ± 0.00 | 7.80 ± 7.09 |  |  |  |
| 3 |  | Amikacin (30) | 25 ± 2.09 | S | 0.83 | 19.83 ± 8.50** | 0.00 ± 0.00 | 0.83 ± 1.33 |  |  |  |
| 4 |  | Kanamycin (30) | 21 ± 0.75 | S | 0.69 | 21.17 ± 8.82*** | 0.00 ± 0.00 | 3.00 ± 2.45 |  |  |  |
| 5 | Fluoroquinolone | Sparfloxacin (5) | 40 ± 0.52 | S | 7.93 | 34.67 ± 15.44*** | 34.50 ± 16.37*** | 46.67 ± 15.41*** | 0.39 | 0.33 | 0.43 |
| 6 |  | Gatifloxacin (5) | 38 ± 0.41 | S | 7.57 | 34.00 ± 13.17*** | 31.40 ± 19.15** | 18.60 ± 12.66** |  |  |  |
| 7 |  | Ofloxacin (5) | 32 ± 1.61 | S | 6.33 | 38.00 ± 17.41*** | 16.17 ± 12.97** | 7.50 ± 7.74 |  |  |  |
| 8 |  | Norfloxacin (10) | 42 ± 1.56 | S | 4.17 | 31.83 ± 16.96** | 3.67 ± 4.23* | 0.00 ± 0.00 |  |  |  |
| 9 |  | Ciprofloxacin (5) | 34 ± 0.63 | S | 6.80 | 7.20 ± 4.15 | 3.50 ± 5.43 | 6.00 ± 8.51 |  |  |  |
| 10 |  | Levofloxacin (5) | 38 ± 1.82 | S | 7.50 | 34.25 ± 3.30*** | 23.40 ± 8.88*** | 29.00 ± 8.19*** |  |  |  |
| 11 |  | Moxifloxacin (5) | 43 ± 0.98 | S | 8.63 | 40.50 ± 18.02*** | 31.00 ± 11.58*** | 35.80 ± 9.39*** |  |  |  |
| 12 | Quinolone (1<sup>st</sup> generation) | Nalidixic acid (30) | 18 ± 0.52 | - | 0.59 | 6.80 ± 9.34 | 3.00 ± 7.35 | 5.83 ± 9.09 | Not applicable |  |  |
| 13 | 3<sup>rd</sup> generation cephalosporin | Ceftriaxone (30) | 30 ± 1.87 | S | 1.01 | 22.83 ± 14.92** | 4.33 ± 4.04* | 0.00 ± 0.00 | -0.32 | -0.90 | -0.65 |
| 14 |  | Cefotaxime (30) | 35 ± 0.55 | - | 1.15 | 17.75 ± 5.74** | 1.00 ± 2.00 | 4.40 ± 5.73 |  |  |  |
| 15 |  | Cefixime (5) | 20 ± 0.52 | - | 3.93 | 20.50 ± 4.68*** | 6.00 ± 5.83* | 8.00 ± 8.67 |  |  |  |
| 16 | Carbapenem | Imipenem (10) | 42 ± 2.61 | - | 4.20 | 7.80 ± 8.50 | 0.00 ± 0.00 | 15.25 ± 2.22*** | Not applicable |  |  |
| 17 | Aminopenicillin | Ampicillin (10) | 31 ± 2.04 | S | 3.08 | 38.33 ± 21.20** | 0.67 ± 1.63 | 21.00 ± 10.23*** |  |  |  |
| 18 | Glycopeptide | Teicoplanin complex (10) | 25 ± 0.52 | S | 2.47 | 31.40 ± 11.87*** | 0.67 ± 1.63 | 6.20 ± 8.84 |  |  |  |
| 19 | Macrolide | Erythromycin (15) | 0 ± 0.00 | R | 0.00 | 21.00 ± 13.08** | 2.00 ± 4.90 | 1.00 ± 2.24 | 0.97 | -0.5 | 0.99 |
| 20 |  | Azithromycin (15) | 0 ± 0.00 | R | 0.00 | 22.50 ± 7.42*** | 0.00 ± 0.00 | 1.50 ± 3.67 |  |  |  |
| 21 |  | Clarithromycin (15) | 11 ± 1.15 | I/R | 0.76 | 28.00 ± 11.96*** | 0.00 ± 0.00 | 8.50 ± 9.01 |  |  |  |
| 22 | Oxazolidinone | Linezolid (30) | 40 ± 0.63 | S | 1.33 | 37.25 ± 3.95*** | 0.00 ± 0.00 | 35.80 ± 11.99*** | Not applicable |  |  |
| 23 | Tetracycline | Doxycycline (hydrochloride) (30) | 40 ± 0.41 | - | 1.34 | 36.50 ± 11.29*** | 35.50 ± 17.72*** | 34.67 ± 13.25*** |  |  |  |
| 24 | Amphenicol | Chloramphenicol (30) | 31 ± 1.67 | S | 1.03 | 56.00 ± 16.24*** | 28.25 ± 8.42*** | 0.00 ± 0.00 |  |  |  |
| 25 | Aminocoumarin | Novobiocin (5) | 38 ± 1.17 | S | 7.63 | 23.40 ± 10.95** | 0.00 ± 0.00 | 7.00 ± 6.78 |  |  |  |
| 26 | Rifamycin | Rifampicin (5) | 41 ± 0.58 | - | 8.27 | 30.50 ± 11.47*** | 27.80 ± 12.97*** | 1.17 ± 1.83 |  |  |  |
| Overall Correlation across all antibiotic classes (r) |  |  |  |  |  |  |  |  | 0.31 | 0.49 | 0.48 |

\*p ≤ 0.05; \*\*p ≤ 0.01; \*\*\*p ≤ 0.001; Red= No activity, Blue = statistically higher than Infection Control and/or at par to the Health Control, Green= Statistically higher than the Health Control; Values shown here are derived from two or more independent assays; # = Clinical and Laboratory Standards Institute (CLSI) (2024); '-'means no interpretation guide is available; Antibiotics whose name is indicated in red fonts are those where there is a mismatch between results of Disc Diffusion Assay and the in vivo assay; those whose name is in **bold** supported not only survival but also fertility in worm population in face of pathogen-challenge.

**Table 2.**
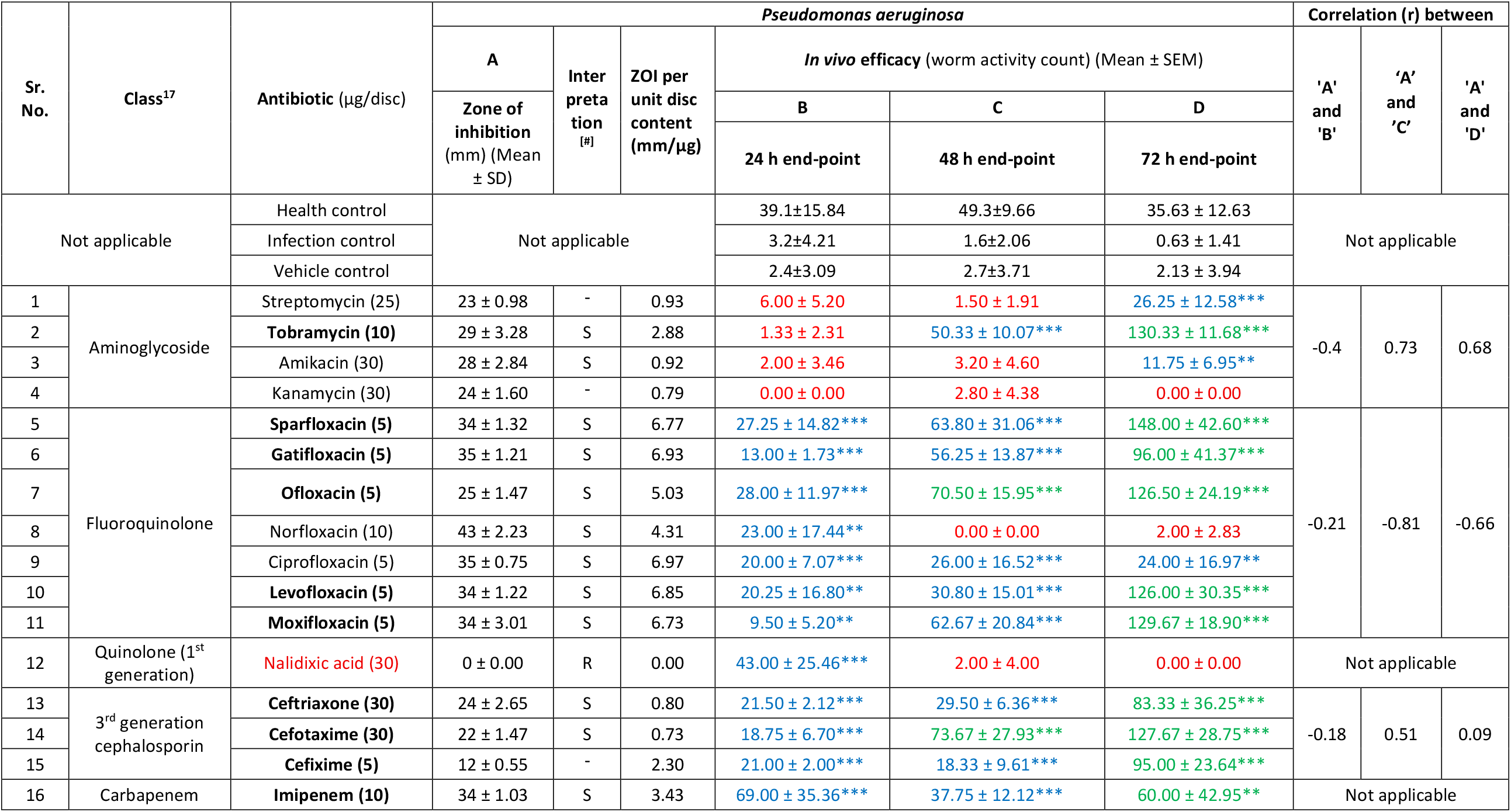
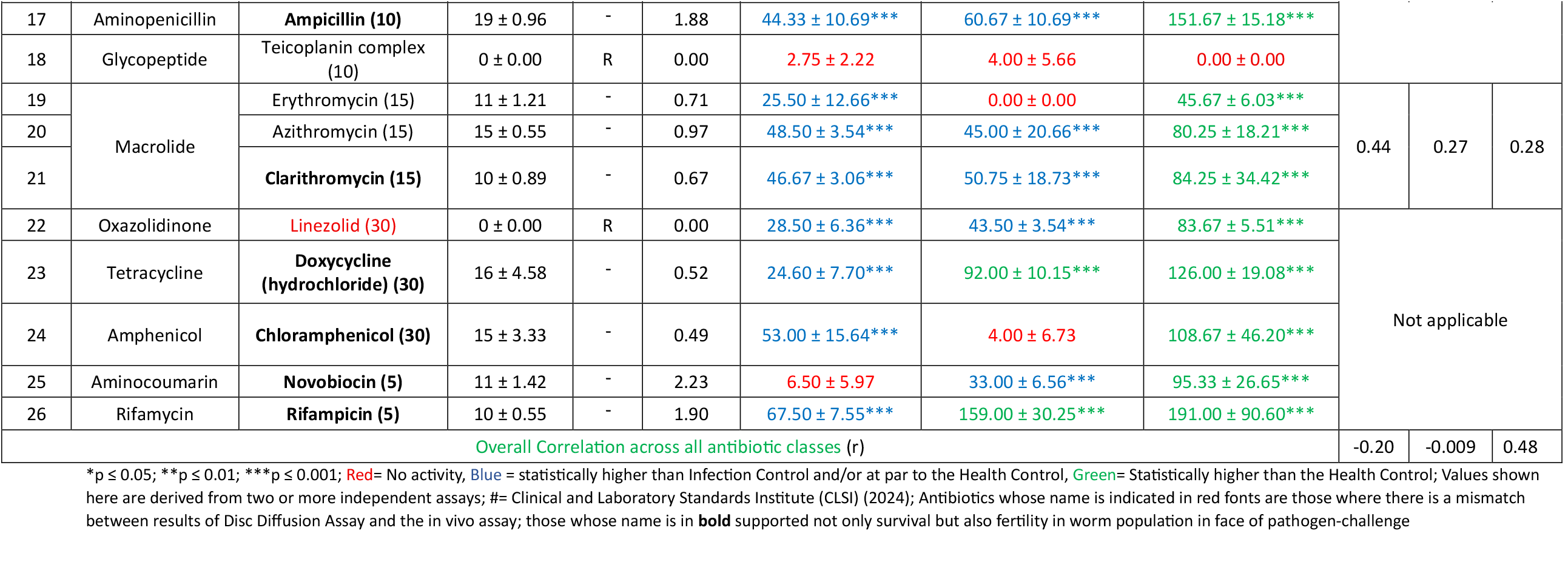
Comparison of *in vitro* and *in vivo* efficacy of antibiotics against *Pseudomonas aeruginosa*.

| Sr. No. | Class<sup>17</sup> | Antibiotic (µg/disc) | *Pseudomonas aeruginosa* |  |  |  |  |  | Correlation (r) between |  |  |
|---|---|---|---|---|---|---|---|---|---|---|---|
|  |  |  | A | Interpretation<sup>[#]</sup> | ZOI per unit disc content (mm/µg) | *In vivo* efficacy (worm activity count) (Mean ± SEM) |  |  | 'A' and 'B' | ‘A’ and ’C’ | 'A' and 'D' |
|  |  |  | Zone of inhibition (mm) (Mean ± SD) |  |  | B | C | D |  |  |  |
|  |  |  |  |  |  | 24 h end-point | 48 h end-point | 72 h end-point |  |  |  |
| Not applicable |  | Health control | Not applicable |  |  | 39.1±15.84 | 49.3±9.66 | 35.63 ± 12.63 | Not applicable |  |  |
|  |  | Infection control |  |  |  | 3.2±4.21 | 1.6±2.06 | 0.63 ± 1.41 |  |  |  |
|  |  | Vehicle control |  |  |  | 2.4±3.09 | 2.7±3.71 | 2.13 ± 3.94 |  |  |  |
| 1 | Aminoglycoside | Streptomycin (25) | 23 ± 0.98 | - | 0.93 | 6.00 ± 5.20 | 1.50 ± 1.91 | 26.25 ± 12.58*** | -0.4 | 0.73 | 0.68 |
| 2 |  | **Tobramycin (10)** | 29 ± 3.28 | S | 2.88 | 1.33 ± 2.31 | 50.33 ± 10.07*** | 130.33 ± 11.68*** |  |  |  |
| 3 |  | Amikacin (30) | 28 ± 2.84 | S | 0.92 | 2.00 ± 3.46 | 3.20 ± 4.60 | 11.75 ± 6.95** |  |  |  |
| 4 |  | Kanamycin (30) | 24 ± 1.60 | - | 0.79 | 0.00 ± 0.00 | 2.80 ± 4.38 | 0.00 ± 0.00 |  |  |  |
| 5 | Fluoroquinolone | **Sparfloxacin (5)** | 34 ± 1.32 | S | 6.77 | 27.25 ± 14.82*** | 63.80 ± 31.06*** | 148.00 ± 42.60*** | -0.21 | -0.81 | -0.66 |
| 6 |  | **Gatifloxacin (5)** | 35 ± 1.21 | S | 6.93 | 13.00 ± 1.73*** | 56.25 ± 13.87*** | 96.00 ± 41.37*** |  |  |  |
| 7 |  | **Ofloxacin (5)** | 25 ± 1.47 | S | 5.03 | 28.00 ± 11.97*** | 70.50 ± 15.95*** | 126.50 ± 24.19*** |  |  |  |
| 8 |  | Norfloxacin (10) | 43 ± 2.23 | S | 4.31 | 23.00 ± 17.44** | 0.00 ± 0.00 | 2.00 ± 2.83 |  |  |  |
| 9 |  | Ciprofloxacin (5) | 35 ± 0.75 | S | 6.97 | 20.00 ± 7.07*** | 26.00 ± 16.52*** | 24.00 ± 16.97** |  |  |  |
| 10 |  | **Levofloxacin (5)** | 34 ± 1.22 | S | 6.85 | 20.25 ± 16.80** | 30.80 ± 15.01*** | 126.00 ± 30.35*** |  |  |  |
| 11 |  | **Moxifloxacin (5)** | 34 ± 3.01 | S | 6.73 | 9.50 ± 5.20** | 62.67 ± 20.84*** | 129.67 ± 18.90*** |  |  |  |
| 12 | Quinolone (1<sup>st</sup> generation) | Nalidixic acid (30) | 0 ± 0.00 | R | 0.00 | 43.00 ± 25.46*** | 2.00 ± 4.00 | 0.00 ± 0.00 | Not applicable |  |  |
| 13 | 3<sup>rd</sup> generation cephalosporin | **Ceftriaxone (30)** | 24 ± 2.65 | S | 0.80 | 21.50 ± 2.12*** | 29.50 ± 6.36*** | 83.33 ± 36.25*** | -0.18 | 0.51 | 0.09 |
| 14 |  | **Cefotaxime (30)** | 22 ± 1.47 | S | 0.73 | 18.75 ± 6.70*** | 73.67 ± 27.93*** | 127.67 ± 28.75*** |  |  |  |
| 15 |  | **Cefixime (5)** | 12 ± 0.55 | - | 2.30 | 21.00 ± 2.00*** | 18.33 ± 9.61*** | 95.00 ± 23.64*** |  |  |  |
| 16 | Carbapenem | **Imipenem (10)** | 34 ± 1.03 | S | 3.43 | 69.00 ± 35.36*** | 37.75 ± 12.12*** | 60.00 ± 42.95** | Not applicable |  |  |
| 17 | Aminopenicillin | **Ampicillin (10)** | 19 ± 0.96 | - | 1.88 | 44.33 ± 10.69*** | 60.67 ± 10.69*** | 151.67 ± 15.18*** |  |  |  |
| 18 | Glycopeptide | Teicoplanin complex (10) | 0 ± 0.00 | R | 0.00 | 2.75 ± 2.22 | 4.00 ± 5.66 | 0.00 ± 0.00 |  |  |  |
| 19 | Macrolide | Erythromycin (15) | 11 ± 1.21 | - | 0.71 | 25.50 ± 12.66*** | 0.00 ± 0.00 | 45.67 ± 6.03*** | 0.44 | 0.27 | 0.28 |
| 20 |  | Azithromycin (15) | 15 ± 0.55 | - | 0.97 | 48.50 ± 3.54*** | 45.00 ± 20.66*** | 80.25 ± 18.21*** |  |  |  |
| 21 |  | **Clarithromycin (15)** | 10 ± 0.89 | - | 0.67 | 46.67 ± 3.06*** | 50.75 ± 18.73*** | 84.25 ± 34.42*** |  |  |  |
| 22 | Oxazolidinone | Linezolid (30) | 0 ± 0.00 | R | 0.00 | 28.50 ± 6.36*** | 43.50 ± 3.54*** | 83.67 ± 5.51*** | Not applicable |  |  |
| 23 | Tetracycline | **Doxycycline (hydrochloride) (30)** | 16 ± 4.58 | - | 0.52 | 24.60 ± 7.70*** | 92.00 ± 10.15*** | 126.00 ± 19.08*** |  |  |  |
| 24 | Amphenicol | **Chloramphenicol (30)** | 15 ± 3.33 | - | 0.49 | 53.00 ± 15.64*** | 4.00 ± 6.73 | 108.67 ± 46.20*** |  |  |  |
| 25 | Aminocoumarin | **Novobiocin (5)** | 11 ± 1.42 | - | 2.23 | 6.50 ± 5.97 | 33.00 ± 6.56*** | 95.33 ± 26.65*** |  |  |  |
| 26 | Rifamycin | **Rifampicin (5)** | 10 ± 0.55 | - | 1.90 | 67.50 ± 7.55*** | 159.00 ± 30.25*** | 191.00 ± 90.60*** |  |  |  |
| Overall Correlation across all antibiotic classes (r) |  |  |  |  |  |  |  |  | -0.20 | -0.009 | 0.48 |

\*p ≤ 0.05; \*\*p ≤ 0.01; \*\*\*p ≤ 0.001; Red= No activity, Blue = statistically higher than Infection Control and/or at par to the Health Control, Green= Statistically higher than the Health Control; Values shown here are derived from two or more independent assays; #= Clinical and Laboratory Standards Institute (CLSI) (2024); Antibiotics whose name is indicated in red fonts are those where there is a mismatch between results of Disc Diffusion Assay and the in vivo assay; those whose name is in **bold** supported not only survival but also fertility in worm population in face of pathogen-challenge

**Table 3.** Comparison of *in vitro* and *in vivo* efficacy of antibiotics against *Clostridioides difficile*.

| Sr. No. | **Class<sup>17</sup>** | **Antibiotic** (µg/disc) | ***Clostridioides difficile*** |  |  |  |  |  | **Correlation (r) between** |  |  |
|---|---|---|---|---|---|---|---|---|---|---|---|
|  |  |  | **A** | **Interpr-etation<sup>#</sup>** | **ZOI per unit disc content (mm/µg)** | ***In vivo* efficacy** (worm activity count) (Mean ± SEM) |  |  | **'A' and 'B'** | **'A' and 'C'** | **'A' and 'D'** |
|  |  |  | **Zone of inhibition** (mm) (Mean ± SD) |  |  | **B** | **C** | **D** |  |  |  |
|  |  |  |  |  |  | **24 h end-point** | **48 h end-point** | **72 end-point** |  |  |  |
| Not applicable |  | Health control | Not applicable |  |  | 29.36±9.73 | 21.77±12.88 | 29.00 ± 7.72 | Not applicable |  |  |
|  |  | Infection control |  |  |  | 2.94±1.18 | 0±0 | 0 ± 0.00 |  |  |  |
|  |  | Vehicle control |  |  |  | 2.88±1.95 | 0±0 | 0 ± 0.00 |  |  |  |
| 1 | Aminoglycoside | Streptomycin (25) | 26 ± 1.17 | S | 1.05 | 0.00 ± 0.00*** | 0.00 ± 0.00 | 0.00 ± 0.00 | -0.49 | NA | -0.62 |
| 2 |  | Tobramycin (10) | 26 ± 1.38 | S | 2.64 | 48.00 ± 16.97 (p=0.06) | 58.50 ± 82.73 | 0.00 ± 0.00 |  |  |  |
| 3 |  | Amikacin (30) | 29 ± 1.06 | S | 0.96 | 0.00 ± 0.00*** | 0.00 ± 0.00 | 0.33 ± 0.82 |  |  |  |
| 4 |  | Kanamycin (30) | 28 ± 0.41 | S | 0.94 | 0.00 ± 0.00*** | 0.00 ± 0.00 | 0.00 ± 0.00 |  |  |  |
| 5 | Fluoroquinolone | **Sparfloxacin (5)** | 30 ± 1.03 | S | 6.07 | 43.5 ± 8.57*** | 18.33 ± 8.02** | 70.80 ± 41.22* | -0.67 | -0.34 | 0.49 |
| 6 |  | Gatifloxacin (5) | 31 ± 0.82 | S | 6.13 | 30.33 ± 21.21 | 50.75 ± 24.66** | 56.33 ± 77.00 |  |  |  |
| 7 |  | Ofloxacin (5) | 27 ± 1.22 | S | 5.35 | 37.83 ± 14.46** | 16.5 ± 12.02 | 60.17 ± 89.45 |  |  |  |
| 8 |  | Norfloxacin (10) | 34 ± 1.78 | S | 3.41 | 25 ± 11.04** | 48.00 ± 68.06 | 64.00 ± 100.40 |  |  |  |
| 9 |  | Ciprofloxacin (5) | 29 ± 1.10 | S | 5.80 | 38.2 ± 17.97** | 78 ± 49.66* | 68.00 ± 104.97 |  |  |  |
| 10 |  | Levofloxacin (5) | 31 ± 1.16 | S | 6.12 | 31.2 ± 12.51** | 0.5 ± 1.00 | 2.17 ± 5.31 |  |  |  |
| 11 |  | Moxifloxacin (5) | 30 ± 0.98 | S | 6.03 | 32.2 ± 8.07*** | 37 ± 22.31* | 28.00 ± 26.98 |  |  |  |
| 12 | Quinolone (1<sup>st</sup> generation) | Nalidixic acid (30) | 22 ± 0.55 | S | 0.72 | 0.00 ± 0.00 | 0.00 ± 0.00 | 0.00 ± 0.00 | Not applicable |  |  |
| 13 | 3<sup>rd</sup> generation cephalosporin | Ceftriaxone (30) | 15 ± 1.13 | S | 0.51 | 34.67 ± 12.50** | 0.00 ± 0.00 | 0.00 ± 0.00 | 0.93 | -0.99 | 0.55 |
| 14 |  | Cefotaxime (30) | 15 ± 0.75 | S | 0.51 | 22.00 ± 3.61*** | 0.00 ± 0.00 | 2.00 ± 4.90 |  |  |  |
| 15 |  | Cefixime (5) | 0 ± 0.00 | R | 0.00 | 0.00 ± 0.00 | 1.50 ± 2.12 | 0.00 ± 0.00 |  |  |  |
| 16 | Carbapenem | Imipenem (10) | 41 ± 0.89 | S | 4.10 | 0.00 ± 0.00 | 0.00 ± 0.00 | 0.00 ± 0.00 | Not applicable |  |  |
| 17 | Aminopenicillin | Ampicillin (10) | 12 ± 0.78 | S | 1.23 | 0.00 ± 0.00 | 0.00 ± 0.00 | 3.83 ± 6.15 |  |  |  |
| 18 | Glycopeptide | Teicoplanin complex (10) | 21 ± 0.82 | S | 2.07 | 13.66 ± 15.62 | 0.00 ± 0.00 | 0.67 ± 1.63 |  |  |  |
| 19 | Macrolide | Erythromycin (15) | 30 ± 2.17 | S | 1.97 | 28.5 ± 8.96** | 20.80 ± 28.91 | 7.00 ± 10.86 | -0.89 | NA | -0.89 |
| 20 |  | Azithromycin (15) | 25 ± 1.21 | S | 1.64 | 43.00 ± 8.49* | 0.00 ± 0.00 | 12.67 ± 19.03 |  |  |  |
| 21 |  | Clarithromycin (15) | 32 ± 1.03 | S | 2.11 | 31.00 ± 10.44** | 0.00 ± 0.00 | 4.33 ± 8.80 |  |  |  |
| 22 | Oxazolidinone | Linezolid (30) | 31 ± 0.82 | S | 1.04 | 15.33 ± 17.61 | 0.00 ± 0.00 | 15.60 ± 24.17 | Not applicable |  |  |
| 23 | Tetracycline | Doxycycline (hydrochloride) (30) | 32 ± 1.63 | S | 1.06 | 40 ± 24.38* | 39.4 ± 19.46** | 50.20 ± 54.11 |  |  |  |
| 24 | Amphenicol | Chloramphenicol (30) | 28 ± 1.41 | S | 0.93 | 25.8 ± 12.73* | 34.60 ± 47.39 | 0.00 ± 0.00 |  |  |  |
| 25 | Aminocoumarin | Novobiocin (5) | 19 ± 1.55 | S | 3.80 | 5.50 ± 0.71 | 0.00 ± 0.00 | 0.00 ± 0.00 |  |  |  |
| 26 | Rifamycin | Rifampicin (5) | 17 ± 1.33 | S | 3.37 | 43 ± 15.85** | 29.80 ± 35.24 | 0.50 ± 1.22 |  |  |  |
| Overall Correlation across all antibiotic classes (r) |  |  |  |  |  |  |  |  | 0.22 | 0.25 | 0.08 |

\*p ≤ 0.05; \*\*p ≤ 0.01; \*\*\*p ≤ 0.001; Red= No activity, Blue = statistically higher than Infection Control and/or at par to the Health Control, Green= Statistically higher than the Health Control; Values shown here are derived from two or more independent assays; <sup>#</sup>Interpretation guide for Antimicrobial Susceptibility Test (AST) through DDA not available, hence we empirically have considered this bacterium susceptible to any antibiotic which could produce an inhibition zone of >10 mm; antibiotics whose name is indicated in red fonts are those where there is a mismatch between results of Disc Diffusion Assay and the in vivo assay; those whose name is in **bold** supported not only survival but also fertility in worm population in face of pathogen-challenge; NA = Not Applicable

## 3. Results and Discussion

Antibiogram of each of the bacterial pathogen was first generated through disc diffusion assay (DDA), wherein bacterial susceptibility was checked against a total of 42 antibiotics (Table S1). Each of the pathogen was also checked for their *in vivo* susceptibility to a panel of 74-75 antibiotics (Table S2). Then further analysis with respect to investigating correlation between DDA and *in vivo* assay was done using those 26 antibiotics which were common between both the assays (Table 1-3).

The present study compared the performance of conventional DDA with an *in vivo Caenorhabditis elegans* infection model for evaluating the efficacy of 26 antibiotics belonging to multiple antimicrobial classes against multi drug resistant *Pseudomonas aeruginosa, Clostridioides difficile*, and methicillin-resistant *Staphylococcus aureus* (MRSA). In the worm assay, antibiotic efficacy was assessed at 24, 48, and 72 h from quantitative worm activity, with infection and vehicle controls providing pathogen- and solvent-associated baselines. The principal observation was that susceptibility inferred from DDA did not consistently predict antibiotic efficacy in the infected model host. This distinction is important because DDA is a standardized antimicrobial susceptibility method based on inhibition of bacterial growth and interpretation according to validated criteria, whereas the *C. elegans* assay measures the capacity of an antibiotic to protect a living host from the consequences of infection.

The overall relationship between Zone of inhibition (ZOI) and *in vivo* worm activity was weak and pathogen dependent. At 24 h, the reported correlations between ZOI and worm activity were *r* = −0.20 for *P. aeruginosa, r* = 0.22 for *C. difficile*, and *r* = 0.31 for MRSA; corresponding 48 h values were −0.009, 0.25, and 0.49, respectively. Thus, none of the three pathogen–host systems showed a strong overall correlation between DDA performance and host-level antibiotic efficacy. The mismatch was particularly evident for aminoglycosides against *C. difficile* and *P. aeruginosa*, macrolides against MRSA, oxazolidinones against *C. difficile* and *P. aeruginosa*, and nalidixic acid across the three pathogens. The detailed dataset illustrates this discordance clearly. For example, against *C. difficile*, imipenem produced the largest inhibition zone (41 mm) but no detectable worm protection at 24 or 48 h, whereas tobramycin, with a substantially smaller ZOI of 26 mm, produced the highest early worm activity (48.0 ± 16.97 at 24 h).

The weak correspondence between the two assays is biologically plausible because DDA and the nematode model interrogate different aspects of antimicrobial action. ZOI reflects bacterial growth inhibition under defined agar conditions and is additionally influenced by diffusion, physicochemical properties, and the amount of antibiotic present on the disk. In contrast, a whole-animal assay incorporates drug availability within the assay environment, pathogen–host interactions, pathogen virulence, and the ability of the compound to preserve host function during infection. *C. elegans* has been established as a useful host model for studying bacterial pathogenesis and for antimicrobial discovery, including the identification of compounds whose activity cannot be predicted solely from bacterial growth inhibition (Ewbank & Zugasti, 2011; Kong et al., 2016). The present findings are therefore consistent with the broader concept that a whole-organism assay can provide biological information that is not captured by a conventional antibiogram.

Several published studies provide useful context for interpreting these findings. Earlier work established that *C. elegans* can reproduce biologically relevant consequences of bacterial infection: fitness costs of antibiotic resistance observed in the nematode were comparable to those measured in mice, supporting its use as a simple host model for studying antibiotic-resistant bacteria (Paulander et al., 2007). Similarly, studies of MRSA demonstrated that different clinical MRSA lineages can exhibit markedly different pathogenicity in *C. elegans*, highlighting the influence of strain-specific host– pathogen interactions on the observed phenotype (Wu et al., 2010). These studies support the premise that antimicrobial performance in a host cannot necessarily be inferred from bacterial susceptibility alone.

An especially relevant comparison is provided by Kong et al. (2014), who developed a liquid-based *C. elegans-S. aureus* model to identify anti-infective agents. Among the positive hits, a substantial proportion improved survival without inhibiting *S. aureus* growth *in vitro*. Specifically, 8 of 28 positive hits had no detectable antibacterial effect in the corresponding *in vitro* assay, while several of these compounds protected worms from MRSA infection. This finding demonstrates that host protection can identify biologically active compounds that conventional bacterial-growth assays may overlook. The present study extends this concept in a different direction; rather than searching for novel compounds without prior antimicrobial activity, it directly compares the efficacy ranking of established antibiotics obtained by DDA with their performance in an infected host. The resulting mismatches therefore suggest that conventional susceptibility measurements may not fully predict host-level antibiotic performance even for established antibiotics.

A contrasting observation was reported by Yao et al. (2022), who evaluated antimicrobial efficacy against carbapenem-resistant *Klebsiella pneumoniae* using a *C. elegans* infection model and found good consistency between nematode-based drug sensitivity results and Kirby–Bauer testing across 10 clinical isolates. Their study therefore demonstrates that strong agreement between an agar-based susceptibility assay and a nematode model is possible under particular experimental conditions. The apparent difference from the present study may reflect important methodological differences. Yao et al. (2022) investigated a single pathogen, employed multiple antibiotic concentrations, and assessed relatively early time points, whereas the present study compared 26 antibiotics across three pathogens at a common concentration of 9.6 ppm and followed host activity over 24-72 h. The present experimental design therefore addresses a broader question of whether DDA can predict the *relative efficacy of diverse antibiotics within a host*, rather than whether an individual antibiotic’s susceptibility classification agrees between two testing systems. Notably, Yao et al. also emphasized the need for larger datasets and additional validation before defining nematode-based breakpoints, which is directly relevant to the interpretation of the present results.

The strength of the DDA-*in vivo* relationship also depended on pathogen, antibiotic class, and assay time. MRSA showed a stronger relationship at 48 h than at 24 h, whereas *P. aeruginosa* showed essentially no correlation at either endpoint. For *C. difficile*, the relationship remained weak overall but became stronger for some later measurements. At the antibiotic-class level, marked differences were also evident. For example, third-generation cephalosporins against *C. difficile* showed a strong positive correlation (*r* = 0.93), whereas several other antibiotic classes exhibited weak or negative associations. Because individual classes contained relatively few antibiotics, these class-specific correlations should be considered exploratory; nevertheless, they indicate that the predictive value of ZOI is not uniform across antibiotic classes. This is an important observation because antibiotics differ substantially in diffusion behaviour, mechanism of action, potency, susceptibility to resistance mechanisms, and interaction with host physiology.

We next examined whether normalization of ZOI to the antibiotic concentration on the disk could improve prediction of *in vivo* efficacy. The ZOI/ antibiotic concentration ratio showed some improvement over raw ZOI in selected pathogen–endpoint comparisons, most notably for *C. difficile*. However, the improvement was not consistent across all three pathogens or across all time points. Thus, the ratio appears to be a potentially useful exploratory parameter for comparing antibiotics tested at different disk loads, but the present dataset does not establish it as a universally superior predictor. This finding is also conceptually consistent with the limitations of diffusion-based measurements, in which observed zone diameters reflect both antimicrobial activity and diffusion characteristics (Lewis, 2024).

The different assays also produced markedly different rankings of antibiotic efficacy. Against *P. aeruginosa*, norfloxacin produced the largest ZOI (43 mm), whereas imipenem and rifampicin produced much higher early worm activity (69.0 ± 35.36 and 67.5 ± 7.55, respectively). Against *C. difficile*, imipenem had the largest ZOI (41 mm), but tobramycin produced the greatest 24 h worm activity. Against MRSA, moxifloxacin produced the largest ZOI (43 mm), whereas chloramphenicol produced the highest 24 h worm activity (56.0 ± 16.24). Therefore, ranking antibiotics according to ZOI, ZOI/concentration, or host activity can yield substantially different conclusions regarding which antibiotic is ‘most effective’. The within-class analysis (Table S3-S5) further supports this finding. For *C. difficile*, for example, tobramycin was statistically distinct from the other aminoglycosides at both 24 and 48 h, whereas streptomycin, amikacin, and kanamycin showed essentially no protective activity.

The host-level phenotype used in the present study is also consistent with the established utility of *C. elegans* for antimicrobial screening. Ewbank & Zugasti, (2011) emphasized that the nematode provides a tractable whole-animal platform for studying antimicrobial activity and host– pathogen interactions, while Kong et al. (2016) highlighted its value for identifying compounds acting through anti-virulence or host-directed mechanisms that may not produce conventional antibacterial phenotypes. The automated measurement of worm activity used here extends this principle by providing a quantitative functional readout of host health rather than relying solely on endpoint mortality. The supplementary videos (A-O) (https://zenodo.org/records/24302220), when looked along with antibiogram (Figure S1-S3), further strengthen this interpretation by providing direct visual evidence of the differences in motility and activity underlying the numerical measurements.

Taken together, the present findings provide evidence against using DDA alone as a surrogate for host-level antibiotic efficacy. However, they do not justify replacing standardized DDA with the *C. elegans* assay for routine clinical antimicrobial susceptibility testing. DDA remains a validated standardized method with established interpretive criteria, whereas the present worm assay has not been calibrated to clinical breakpoints or therapeutic outcomes. Moreover, the study itself has important limitations: all antibiotics were assessed at a single concentration of 9.6 ppm, and differences between the agar-based DDA environment and the M9-based environment used for the *in vivo* assay may have contributed to the relatively poor correspondence.

Instead, the present findings support a complementary role for the nematode model and provide a rationale for further development of a 24 h *C. elegans* antibiotic-efficacy assay as a host-level screening tool. Such an assay could potentially be particularly valuable for prioritizing antibiotics for further therapeutic investigation, because it asks a functionally different question from DDA: not simply whether bacterial growth is inhibited, but whether the antibiotic can protect an infected host. This interpretation is compatible with previous literature showing that *C. elegans* can identify anti-infective activity that is not evident from conventional antimicrobial assays (Kong et al., 2014; Kong et al., 2016), while also remaining consistent with studies demonstrating that concordance with Kirby– Bauer testing can occur under other experimental conditions (Yao et al., 2022).

Several limitations of the present study should be considered while interpreting its findings. First, only one strain of each bacterial pathogen was investigated; therefore, the observed discrepancies between disc diffusion assay (DDA) results and host-protective antibiotic activity cannot necessarily be generalized to other strains or clinical isolates. Second, all antibiotics were evaluated at a single concentration (9.6 µg/mL) in the *C. elegans* infection assay. Since antibiotics differ considerably in their molecular weights, pharmacological properties and effective concentration ranges, a common mass concentration may not represent equivalent antimicrobial exposure across different compounds. Third, DDA and the nematode infection assay were performed under fundamentally different experimental conditions, with the former evaluating bacterial growth inhibition in an agar-based environment and the latter incorporating host-pathogen interactions in a liquid-based system. These differences may have contributed to the observed discrepancies between the two approaches. Furthermore, minimum inhibitory concentration (MIC) measurements were not performed, which could have provided an additional basis for comparison with the host-protective activity observed in the nematode model. Finally, the 72 h worm activity measurements require cautious interpretation because progeny production in some antibiotic-treated groups could contribute to the recorded activity counts, making these measurements reflective of both parental worm and progeny activity rather than the condition of the original worm population alone. Overall, the present data indicate that DDA-derived zone of inhibition (ZOI), ZOI normalized to antibiotic disc content, and *in vivo* worm activity represent related but distinct measures of antibiotic performance. The generally weak correlations, pathogen- and antibiotic-class-dependent relationships, and substantial differences in antibiotic rankings demonstrate that an antibiotic producing the largest inhibition zone is not necessarily the one providing the greatest protection to an infected host. The *C. elegans* model therefore appears most appropriately positioned not as an immediate replacement for DDA, but as a complementary whole-organism screening platform capable of providing functional information that may not be captured by conventional susceptibility testing. Further investigations incorporating multiple clinical isolates, dose-response assessments, broth microdilution/MIC measurements, pharmacologically relevant exposure ranges, independent viability and reproductive assessments, and mammalian validation will be necessary to establish the broader applicability and predictive value of this approach in antimicrobial drug discovery and evaluation.

## Supporting information

Supplementary Tables, Figures, and Legend for Videos

## Acknowledgement

Authors thank Nirma University for access to laboratory infrastructure. MV acknowledges Gujarat Government’s support in form of stipend under the SHODH scheme. Antibiotic library used for *in vivo* assays was provided by CC4CARB, which has been funded with Federal funds from the National Institute of Allergy and Infectious Disease, National Institutes of Health, Department of Health and Human Services, under Contract No. 75N93021C00002.

## Supplementary Material

Table S1: Results of Disc Diffusion Assay of all available antibiotic discs against three pathogens Table S2 (a-c): Results of *in vivo* anti-pathogenic assay with all the 72 antibiotics against three pathogens Table S3-S5: Within class comparison of antibiotic efficacy Figures S1-S3: Representative photos of the disc diffusion assays Appendix A: Legend to supplementary videos Videos A-O: Supplementary videos pertaining to anti-pathogenic assays (https://zenodo.org/records/24302220)

## Notes

### Competing Interest Statement

The authors have declared no competing interest.

