## Supplementary Tables, Figures, and Legend for Videos for "How good a predictor the *in vitro* antibiogram is, of *in vivo* antibiotic efficacy? An *in vitro* - *in vivo* integrated approach"

**Table S1. Results of Disc Diffusion Assay of all available antibiotic discs against three pathogens**

| Sr. No. | Class | Antibiotic | Concentration (µg/disc) | Diameter of Zone of Inhibition (mm) (Mean ± SD) | | | | | |
|---|---|---|---|---|---|---|---|---|---|
| | | | | **MRSA** | **Interpretation** | ***P. aeruginosa*** | **Interpretation** | ***Cl. difficile*** | **Interpretation** |
| 1 | Aminoglycoside | Streptomycin | 25 | 29.00 ± 0.89 | S | 23.17 ± 0.98 | - | 26.17 ± 1.17 | S |
| 2 | | Gentamicin | 10 | 31.28 ± 1.27 | S | 26.11 ± 2.68 | S | 28.00 ± 0.91 | S |
| 3 | | Tobramycin | 10 | 14.83 ± 3.24 | I/R | 28.75 ± 3.28 | S | 26.42 ± 1.38 | S |
| 4 | | Amikacin | 30 | 25.00 ± 2.09 | S | 27.50 ± 2.84 | S | 28.79 ± 1.06 | S |
| 5 | | Kanamycin | 30 | 20.83 ± 0.75 | S | 23.83 ± 1.60 | - | 28.17 ± 0.41 | S |
| 6 | Fluoroquinolone | Sparfloxacin | 5 | 39.67 ± 0.52 | S | 33.83 ± 1.32 | S | 30.33 ± 1.03 | S |
| 7 | | Gatifloxacin | 5 | 37.83 ± 0.41 | S | 34.67 ± 1.21 | S | 30.67 ± 0.82 | S |
| 8 | | Ofloxacin | 5 | 31.67 ± 1.61 | S | 25.17 ± 1.47 | S | 26.75 ± 1.22 | S |
| 9 | | Norfloxacin | 10 | 41.67 ± 1.56 | S | 43.08 ± 2.23 | S | 34.08 ± 1.78 | S |
| 10 | | Ciprofloxacin | 5 | 34.00 ± 0.63 | S | 34.83 ± 0.75 | S | 29.00 ± 1.10 | S |
| 11 | | Levofloxacin | 5 | 37.50 ± 1.82 | S | 34.25 ± 1.22 | S | 30.58 ± 1.16 | S |
| 12 | | Moxifloxacin | 5 | 43.17 ± 0.98 | S | 33.67 ± 3.01 | S | 30.17 ± 0.98 | S |
| 13 | Quinolone (1st generation) | Nalidixic acid | 30 | 17.67 ± 0.52 | - | 0.00 ± 0.00 | R | 21.50 ± 0.55 | S |
| 14 | Cephalosporin | Cefpodoxime | 10 | 20.42 ± 3.23 | S | 0.00 ± 0.00 | R | 0.00 ± 0.00 | R |
| 15 | | Ceftriaxone | 30 | 30.28 ± 1.87 | S | 24.11 ± 2.65 | S | 15.28 ± 1.13 | S |
| 16 | | Cefotaxime | 30 | 34.50 ± 0.55 | - | 21.83 ± 1.47 | - | 15.17 ± 0.75 | S |
| 17 | | Cefixime | 5 | 19.67 ± 0.52 | - | 11.50 ± 0.55 | - | 0.00 ± 0.00 | R |
| 18 | | Cefepime | 30 | 33.17 ± 2.86 | - | 31.67 ± 2.25 | - | 18.83 ± 1.33 | S |
| 19 | | Cephalothin | 30 | 42.00 ± 2.61 | - | 0.00 ± 0.00 | R | 11.83 ± 0.75 | S |
| 20 | Carbapenem | Imipenem | 10 | 50.50 ± 1.23 | - | 34.33 ± 1.03 | S | 41.00 ± 0.89 | S |
| 21 | Penicillin | Penicillin-G | 10 units | 36.00 ± 6.23 | - | 0.00 ± 0.00 | - | 13.17 ± 1.33 | S |
| 22 | Aminopenicillin | Ampicillin | 10 | 30.83 ± 2.04 | - | 18.75 ± 0.96 | - | 12.33 ± 0.78 | S |

| Sr. No. | Class | Antibiotic | Concentration (µg/disc) | Diameter of Zone of Inhibition (mm) (Mean ± SD) | | | | | |
|---|---|---|---|---|---|---|---|---|---|
| | | | | **MRSA** | **Interpretation** | ***P. aeruginosa*** | **Interpretation** | ***Cl. difficile*** | **Interpretation** |
| 23 | Penicillinase-resistant penicillin | Methicillin | 5 | 34.33 ± 0.58 | - | 0.00 ± 0.00 | R | 0.00 ± 0.00 | R |
| 24 | Penicillinase-resistant penicillin | Oxacillin | 1 | 29.67 ± 12.08 | - | 0.00 ± 0.00 | R | 0.00 ± 0.00 | R |
| 25 | Antipseudomonal penicillin | Ticarcillin | 75 | 40.17 ± 0.75 | - | 31.17 ± 3.87 | S | 16.50 ± 2.43 | S |
| 26 | β-lactam + β-lactamase inhibitor | Augmentin | 30 | 31.50 ± 0.55 | S | 0.00 ± 0.00 | R | 14.67 ± 1.03 | S |
| 27 | | Amoxyclav | 30 | 28.47 ± 5.04 | - | 0.00 ± 0.00 | R | 12.22 ± 1.52 | S |
| 28 | Glycopeptide | Vancomycin | 30 | 24.17 ± 1.64 | - | 9.75 ± 0.87 | - | 23.42 ± 0.67 | S |
| 29 | | Teicoplanin | 10 | 24.67 ± 0.52 | - | 0.00 ± 0.00 | R | 20.67 ± 0.82 | S |
| 30 | Macrolide | Erythromycin | 15 | 0.00 ± 0.00 | R | 10.67 ± 1.21 | - | 29.50 ± 2.17 | S |
| 31 | | Azithromycin | 15 | 0.00 ± 0.00 | R | 14.50 ± 0.55 | - | 24.67 ± 1.21 | S |
| 32 | | Clarithromycin | 15 | 5.67 ± 1.16 | - | 10.00 ± 0.89 | - | 31.67 ± 1.03 | S |
| 33 | Lincosamide | Clindamycin | 2 | 0.00 ± 0.00 | R | 0.00 ± 0.00 | R | 28.08 ± 1.24 | S |
| 34 | Oxazolidinone | Linezolid | 30 | 40.00 ± 0.63 | - | 0.00 ± 0.00 | R | 31.33 ± 0.82 | S |
| 35 | Tetracycline | Tetracycline | 30 | 40.17 ± 0.72 | - | 15.50 ± 4.58 | - | 28.08 ± 1.56 | S |
| 36 | | Doxycycline (hydrochloride) | 30 | 40.17 ± 0.41 | - | 14.67 ± 3.33 | - | 31.67 ± 1.63 | S |
| 37 | Amphenicol | Chloramphenicol | 30 | 30.78 ± 1.67 | - | 11.17 ± 1.42 | - | 28.00 ± 1.41 | S |
| 38 | Folate pathway inhibitor (sulfonamide combination) | Co-trimoxazole | 25 | 40.33 ± 0.49 | S | 18.92 ± 3.68 | - | 28.33 ± 0.98 | S |
| 39 | Polymyxin | Colistin | 10 | 12.00 ± 0.63 | - | 17.83 ± 1.47 | S | 0.00 ± 0.00 | R |
| 40 | Nitrofuran | Nitrofurantoin | 300 | 20.33 ± 0.52 | - | 0.00 ± 0.00 | R | 21.33 ± 2.66 | S |
| 41 | Aminocoumarin | Novobiocin | 5 | 38.17 ± 1.17 | - | 0.00 ± 0.00 | R | 19.00 ± 1.55 | S |
| 42 | Rifamycin | Rifampicin | 5 | 41.33 ± 0.58 | - | 9.50 ± 0.55 | - | 16.83 ± 1.33 | S |
| Number of antibiotics which could inhibit the bacterial growth (inhibition zone ≥ 10 mm) | | | | 38 | | 29 | | 37 | |

'-': an interpretive guide is not available

**Table S2 (a). Results of *in vivo* anti-pathogenic assay with all the 74 antibiotics against methicillin-resistant *Staphylococcus aureus***

| Sr. No. | Class | Antibiotic | Methicillin-resistant *Staphylococcus aureus* *In vivo* efficacy (worm activity count) (Mean ± SEM) | | |
|---|---|---|---|---|---|
| | | | 24 h end-point | 48 h end-point | 72 h end-point |
| 1 | Aminoglycoside | Amikacin | 19.83 ± 8.50** | 0.00 ± 0.00 | 0.83 ± 1.33 |
| 2 | | Kanamycin | 21.17 ± 8.82** | 0.00 ± 0.00 | 3.00 ± 2.45 |
| 3 | | Streptomycin | 22.67 ± 6.11*** | 0.00 ± 0.00 | 11.80 ± 8.26** |
| 4 | | Tobramycin | 20.67 ± 10.95** | 0.00 ± 0.00 | 7.80 ± 7.09 |
| 5 | Tetracycline | Omadacycline | 10.80 ± 5.93 | 3.00 ± 4.69 | 7.80 ± 4.32** |
| 6 | Amphenicol | Chloramphenicol | 56.00 ± 16.24*** | 28.25 ± 8.42*** | 0.00 ± 0.00 |
| 7 | | Florfenicol | 37.67 ± 15.60*** | 46.33 ± 9.45*** | 14.50 ± 8.89** |
| 8 | | Thiamphenicol | 17.50 ± 5.39*** | 0.00 ± 0.00 | 0.00 ± 0.00 |
| 9 | Antifolate / diaminopyrimidine | Trimethoprim | 27.00 ± 11.94** | 1.67 ± 2.66 | 2.00 ± 3.35 |
| 10 | | Iclaprim | 38.17 ± 11.82*** | 0.00 ± 0.00 | 1.33 ± 2.31 |
| 11 | Antimycobacterial – ethambutol class | Ethambutol | 3.20 ± 4.60 | 0.00 ± 0.00 | 6.20 ± 5.12 |
| 12 | Antimycobacterial – hydrazide | Isoniazid | 6.75 ± 8.42 | 0.00 ± 0.00 | 9.17 ± 11.63 |
| 13 | Antimycobacterial – pyrazinamide class | Pyrazinamide | 4.50 ± 9.00 | 4.83 ± 6.34 | 2.60 ± 2.41 |
| 14 | Bisphenol / antiseptic; FabI inhibitor | Triclosan | 28.00 ± 12.57** | 2.80 ± 4.38 | 0.00 ± 0.00 |
| 15 | Boron-containing antimicrobial / leucyl-tRNA synthetase inhibitor | Epetraborole | 37.00 ± 18.38*** | 30.50 ± 16.31*** | 34.33 ± 20.70** |
| 16 | Coumarin antibiotic | Novobiocin | 23.40 ± 10.95** | 0.00 ± 0.00 | 7.00 ± 6.78 |
| 17 | Cyclic lipopeptide / biosurfactant | Surfactin | 10.60 ± 9.04 | 1.67 ± 4.08 | 1.83 ± 2.23 |
| 18 | Diarylpyridine | Bedaquiline | 28.25 ± 7.59*** | 0.00 ± 0.00 | 12.00 ± 20.20 |
| 19 | Experimental antimicrobial compound | U105920 | 24.17 ± 9.28*** | 0.00 ± 0.00 | 2.17 ± 2.56 |
| 20 | | Az-1 | 0.50 ± 1.00 | 0.00 ± 0.00 | 4.25 ± 5.06 |

| Sr. No. | Class | Antibiotic | Methicillin-resistant *Staphylococcus aureus* | | |
|---|---|---|---|---|---|
| | | | ***In vivo* efficacy** (worm activity count) (Mean ± SEM) | | |
| | | | **24 h end-point** | **48 h end-point** | **72 h end-point** |
| 21 | Experimental LpxC inhibitor | Chir-090 | 20.40 ± 14.19* | 0.00 ± 0.00 | 5.60 ± 5.37 |
| 22 | | pf-5081090 | 27.67 ± 7.17*** | 0.60 ± 1.34 | 1.67 ± 3.20 |
| 23 | Fluoroquinolone | Delafloxacin | 49.00 ± 12.73*** | 39.50 ± 10.52*** | 38.00 ± 12.29*** |
| 24 | | Gatifloxacin | 34.00 ± 13.17*** | 31.40 ± 19.15** | 18.60 ± 12.66** |
| 25 | | levofloxacin | 34.25 ± 3.30*** | 23.40 ± 8.88*** | 29.00 ± 8.19*** |
| 26 | | Lomefloxacin | 26.75 ± 11.81** | 2.20 ± 3.49 | 0.80 ± 1.79 |
| 27 | | Moxifloxacin | 40.50 ± 18.02*** | 31.00 ± 11.58*** | 35.80 ± 9.39*** |
| 28 | | Norfloxacin | 31.83 ± 16.96** | 3.67 ± 4.23* | 0.00 ± 0.00 |
| 29 | | Ofloxacin | 38.00 ± 17.41*** | 16.17 ± 12.97** | 7.50 ± 7.74 |
| 30 | | Sparfloxacin | 34.67 ± 15.44*** | 34.50 ± 16.37*** | 46.67 ± 15.41*** |
| 31 | | Ciprofloxacin | 7.20 ± 4.15 | 3.50 ± 5.43 | 6.00 ± 8.51 |
| 32 | Glycopeptide | Teicoplanin complex | 31.40 ± 11.87*** | 0.67 ± 1.63 | 6.20 ± 8.84 |
| 33 | Glycopeptide/lipoglycopeptide-related | Ramoplanin complex | 29.00 ± 12.84*** | 0.00 ± 0.00 | 4.00 ± 4.62 |
| 34 | Glycylcycline / tetracycline | tigecycline | 9.20 ± 13.08 | 0.00 ± 0.00 | 2.00 ± 2.83 |
| 35 | Ketolide | Telithromycin | 31.80 ± 12.56*** | 1.00 ± 2.45 | 7.40 ± 4.98* |
| 36 | Lipopeptide | Daptomycin | 32.50 ± 13.13*** | 0.00 ± 0.00 | 6.17 ± 3.60** |
| 37 | Macrocyclic peptide | Murepavidin | 24.33 ± 10.97** | 0.00 ± 0.00 | 1.00 ± 2.24 |
| 38 | Macrolide | Clarithromycin | 28.00 ± 11.96*** | 0.00 ± 0.00 | 8.50 ± 9.01 |
| 39 | | Erythromycin | 21.00 ± 13.08** | 2.00 ± 4.90 | 1.00 ± 2.24 |
| 40 | Macrolide – azalide | Azithromycin | 22.50 ± 7.42*** | 0.00 ± 0.00 | 1.50 ± 3.67 |
| 41 | Nitroimidazole antimycobacterial | Pretomanid | 1.50 ± 3.00 | 0.00 ± 0.00 | 8.00 ± 12.33 |
| 42 | Oxazolidinone | Linezolid | 37.25 ± 3.95*** | 0.00 ± 0.00 | 35.80 ± 11.99*** |
| 43 | | Radezolid | 26.60 ± 18.31** | 0.00 ± 0.00 | 1.25 ± 2.50 |
| 44 | | Sutezolid | 11.67 ± 4.93* | 0.00 ± 0.00 | 33.50 ± 4.95*** |
| 45 | | Tedizolid | 20.17 ± 11.69** | 0.83 ± 2.04 | 29.40 ± 13.78*** |
| 46 | Pleuromutilin | Azamulin | 27.40 ± 9.40*** | 0.00 ± 0.00 | 10.00 ± 12.07 |
| 47 | | Lefamulin | 28.67 ± 10.76*** | 0.20 ± 0.45 | 4.67 ± 5.47 |
| 48 | | Retapamulin | 20.67 ± 10.73** | 1.33 ± 2.42 | 4.50 ± 5.45 |
| 49 | | Tiamulin | 30.67 ± 11.31*** | 3.17 ± 4.31 | 5.40 ± 4.10 |

| Sr. No. | Class | Antibiotic | Methicillin-resistant *Staphylococcus aureus* | | |
|---|---|---|---|---|---|
| | | | ***In vivo* efficacy** (worm activity count) (Mean ± SEM) | | |
| | | | **24 h end-point** | **48 h end-point** | **72 h end-point** |
| 50 | | Valnemulin | 37.40 ± 15.40*** | 0.00 ± 0.00 | 2.80 ± 3.83 |
| 51 | Polymyxin / cyclic lipopeptide | Polymyxin b | 4.60 ± 6.39 | 0.67 ± 1.63 | 1.33 ± 3.27 |
| 52 | Quinolone | Cinoxacin | 20.50 ± 7.33*** | 1.33 ± 2.80 | 24.50 ± 4.80*** |
| 53 | | nalidixic Acid | 6.80 ± 9.34 | 3.00 ± 7.35 | 5.83 ± 9.09 |
| 54 | Rifamycin | Rifampicin | 30.50 ± 11.47*** | 27.80 ± 12.97*** | 1.17 ± 1.83 |
| 55 | Spiropyrimidinetrione | Zoliflodacin | 21.50 ± 9.57** | 0.00 ± 0.00 | 0.00 ± 0.00 |
| 56 | Streptogramin | Virginiamycin complex | 21.50 ± 9.11** | 0.83 ± 2.04 | 10.00 ± 10.71 |
| 57 | Sulfonamide | Sulfonilamide | 23.50 ± 9.15*** | 0.00 ± 0.00 | 10.25 ± 9.67* |
| 58 | Sulfonamide / sulfones | Dapsone | 12.00 ± 2.94** | 0.67 ± 1.03 | 2.17 ± 3.71 |
| 59 | Tetracycline | Doxycycline Hydrochloride | 36.50 ± 11.29*** | 35.50 ± 17.72*** | 34.67 ± 13.25*** |
| 60 | | Lymecycline | 1.80 ± 2.49 | 0.00 ± 0.00 | 10.00 ± 11.82 |
| 61 | | Meclocycline | 7.20 ± 8.32 | 4.00 ± 6.69 | 5.25 ± 5.56 |
| 62 | | Oxytetracycline | 4.60 ± 7.80 | 0.67 ± 1.63 | 9.83 ± 9.52* |
| 63 | | Rolitetracycline | 13.33 ± 8.14* | 0.00 ± 0.00 | 15.00 ± 15.00* |
| 64 | Thiopeptide antibiotic | Nosiheptide | 32.17 ± 9.37*** | 2.33 ± 4.08 | 16.00 ± 4.64*** |
| 65 | β-lactam – aminopenicillin | Amoxicillin | 28.00 ± 7.62*** | 1.33 ± 3.27 | 4.00 ± 9.80 |
| 66 | | Ampicillin | 38.33 ± 21.20** | 0.67 ± 1.63 | 21.00 ± 10.23*** |
| 67 | β-lactam – carbapenem | Meropenem | 18.80 ± 7.60** | 0.00 ± 0.00 | 7.00 ± 7.28 |
| 68 | | Imipenem | 7.80 ± 8.50 | 0.00 ± 0.00 | 15.25 ± 2.22*** |
| 69 | Cephalosporin | Cefadroxil | 20.00 ± 9.06** | 0.00 ± 0.00 | 4.00 ± 4.43 |
| 70 | | Cefaclor | 18.83 ± 13.33* | 0.00 ± 0.00 | 2.00 ± 2.45 |
| 71 | | Cefixime | 20.50 ± 4.68*** | 6.00 ± 5.83* | 8.00 ± 8.67 |
| 72 | | Cefotaxime | 17.75 ± 5.74** | 1.00 ± 2.00 | 4.40 ± 5.73 |
| 73 | | Ceftazidine | 13.67 ± 6.25** | 3.00 ± 2.83* | 2.17 ± 5.31 |
| 74 | | Ceftriaxone | 22.83 ± 14.92** | 4.33 ± 4.04* | 0.00 ± 0.00 |

None of the antibiotics, except triclosan, showed toxicity towards worms at the tested concentration till 72 h.

**Table S2 (b). Results of *in vivo* anti-pathogenic assay with all the 75 antibiotics against *Pseudomonas aeruginosa***

| Sr. No. | Class | Antibiotic | *Pseudomonas aeruginosa* *In vivo* efficacy (worm activity count) (Mean ± SEM) | | |
|---|---|---|---|---|---|
| | | | **24 h end-point** | **48 h end-point** | **72 h end-point** |
| 1 | Aminoglycoside | Amikacin | 2.00 ± 3.46 | 3.20 ± 4.60 | 11.75 ± 6.95** |
| 2 | | Kanamycin | 0.00 ± 0.00 | 2.80 ± 4.38 | 0.00 ± 0.00 |
| 3 | | Streptomycin | 6.00 ± 5.20 | 1.50 ± 1.91 | 26.25 ± 12.58*** |
| 4 | | Tobramycin | 1.33 ± 2.31 | 50.33 ± 10.07*** | 130.33 ± 11.68*** |
| 5 | Aminomethylcycline / tetracycline | Omadacycline | 23.67 ± 15.04*** | 66.67 ± 35.85*** | 87.67 ± 18.01*** |
| 6 | Amphenicol | Chloramphenicol | 53.00 ± 15.64*** | 4.00 ± 6.73 | 108.67 ± 46.20*** |
| 7 | | Florfenicol | 58.33 ± 13.20*** | 75.25 ± 17.58*** | 135.75 ± 39.14*** |
| 8 | | Thiamphenicol | 21.50 ± 5.20*** | 3.80 ± 3.19 | 0.00 ± 0.00 |
| 9 | Antifolate / diaminopyrimidine | Trimethoprim | 23.00 ± 11.53*** | 1.00 ± 2.00 | 0.00 ± 0.00 |
| 10 | | Iclaprim | 17.33 ± 2.08*** | 49.00 ± 24.43*** | 115.33 ± 42.36*** |
| 11 | Antimycobacterial | Ethambutol | 22.75 ± 10.72*** | 23.67 ± 14.15*** | 60.00 ± 26.08*** |
| 12 | | Pyrazinamide | 16.25 ± 5.56*** | 62.67 ± 10.02*** | 66.50 ± 37.01*** |
| 13 | Antimycobacterial / isonicotinic acid hydrazide | Isoniazid | 8.00 ± 3.27** | 55.00 ± 35.36*** | 80.00 ± 51.45*** |
| 14 | Boron-containing antimicrobial / leucyl-tRNA synthetase inhibitor | Epetraborole | 21.20 ± 3.11*** | 64.25 ± 17.46*** | 74.25 ± 50.43*** |
| 15 | Coumarin antibiotic | Novobiocin | 6.50 ± 5.97 | 33.00 ± 6.56*** | 95.33 ± 26.65*** |
| 16 | Cyclic lipopeptide | Daptomycin | 34.67 ± 16.29*** | 36.75 ± 23.50*** | 77.67 ± 33.01*** |
| 17 | Cyclic lipopeptide / biosurfactant | Surfactin | 2.67 ± 4.62 | 5.60 ± 7.70 | 1.00 ± 1.73 |
| 18 | Diarylquinoline / diarylpyridine | Bedaquiline | 14.33 ± 2.08*** | 34.33 ± 5.86*** | 25.75 ± 22.84** |
| 19 | Experimental antimicrobial | U105920 | 1.33 ± 2.31 | 0.00 ± 0.00 | 0.00 ± 0.00 |
| 20 | | AZ-1 | 25.67 ± 18.18*** | 38.50 ± 13.44*** | 50.50 ± 29.51*** |
| 21 | Fluoroquinolone | Delafloxacin | 41.00 ± 15.59*** | 52.25 ± 15.59*** | 135.25 ± 18.43*** |
| 22 | | Gatifloxacin | 13.00 ± 1.73*** | 56.25 ± 13.87*** | 96.00 ± 41.37*** |
| 23 | | Levofloxacin | 20.25 ± 16.80** | 30.80 ± 15.01*** | 126.00 ± 30.35*** |
| 24 | | Lomefloxacin | 16.33 ± 8.39*** | 46.75 ± 9.78*** | 81.00 ± 27.07*** |
| 25 | | Moxifloxacin | 9.50 ± 5.20** | 62.67 ± 20.84*** | 129.67 ± 18.90*** |
| 26 | | Norfloxacin | 23.00 ± 17.44** | 0.00 ± 0.00 | 2.00 ± 2.83 |
| 27 | | Ofloxacin | 28.00 ± 11.97*** | 70.50 ± 15.95*** | 126.50 ± 24.19*** |

| Sr. No. | Class | Antibiotic | *Pseudomonas aeruginosa* | | |
|---|---|---|---|---|---|
| | | | ***In vivo* efficacy** (worm activity count) (Mean ± SEM) | | |
| | | | **24 h end-point** | **48 h end-point** | **72 h end-point** |
| 28 | | Sparfloxacin | 27.25 ± 14.82*** | 63.80 ± 31.06*** | 148.00 ± 42.60*** |
| 29 | | Trovafloxacin | 25.50 ± 7.78*** | 30.00 ± 2.83*** | 42.50 ± 3.54*** |
| 30 | | Ciprofloxacin | 20.00 ± 7.07*** | 26.00 ± 16.52*** | 24.00 ± 16.97** |
| 31 | Glycopeptide | Teicoplanin complex | 2.75 ± 2.22 | 4.00 ± 5.66 | 0.00 ± 0.00 |
| 32 | Glycopeptide-related / lipoglycodepsipeptide | Ramoplanin complex | 0.00 ± 0.00 | 2.25 ± 4.50 | 0.00 ± 0.00 |
| 33 | Glycylcycline / tetracycline | Tigecycline | 9.50 ± 8.19* | 16.00 ± 5.66*** | 14.50 ± 14.20* |
| 34 | Ketolide | Telithromycin | 45.25 ± 17.99*** | 23.75 ± 11.15*** | 71.00 ± 31.26*** |
| 35 | LpxC inhibitor | CHIR-090 | 46.67 ± 18.50*** | 0.00 ± 0.00 | 0.00 ± 0.00 |
| 36 | | PF-5081090 | 38.33 ± 5.51*** | 50.75 ± 15.69*** | 84.75 ± 56.26*** |
| 37 | Macrocyclic peptide | Murepavidin | 21.50 ± 2.12*** | 9.25 ± 4.86** | 1.00 ± 2.00 |
| 38 | Macrolide | Clarithromycin | 46.67 ± 3.06*** | 50.75 ± 18.73*** | 84.25 ± 34.42*** |
| 39 | | Erythromycin | 25.50 ± 12.66*** | 0.00 ± 0.00 | 45.67 ± 6.03*** |
| 40 | Macrolide / azalide | Azithromycin | 48.50 ± 3.54*** | 45.00 ± 20.66*** | 80.25 ± 18.21*** |
| 41 | Nitroimidazole antimycobacterial | Pretomanid | 47.33 ± 11.68*** | 133.00 ± 101.82*** | 73.33 ± 26.56*** |
| 42 | Oxazolidinone | Linezolid | 28.50 ± 6.36*** | 43.50 ± 3.54*** | 83.67 ± 5.51*** |
| 43 | | Radezolid | 21.00 ± 9.54*** | 3.00 ± 3.83 | 2.00 ± 2.83 |
| 44 | | Sutezolid | 4.67 ± 0.58 | 0.00 ± 0.00 | 0.00 ± 0.00 |
| 45 | | Tedizolid | 29.33 ± 5.51*** | 3.00 ± 3.83 | 31.67 ± 16.50*** |
| 46 | Pleuromutilin | Azamulin | 31.50 ± 2.12*** | 25.33 ± 0.58*** | 73.50 ± 17.68*** |
| 47 | | Lefamulin | 12.67 ± 9.02** | 24.00 ± 19.16** | 42.75 ± 16.72*** |
| 48 | | Retapamulin | 0.00 ± 0.00 | 0.67 ± 1.15 | 2.00 ± 3.46 |
| 49 | | Tiamulin | 33.67 ± 5.51*** | 28.33 ± 0.58*** | 60.00 ± 17.78*** |
| 50 | | Valnemulin | 29.80 ± 5.93*** | 41.25 ± 15.39*** | 90.67 ± 34.78*** |
| 51 | Polymyxin / cyclic lipopeptide | Polymyxin B | 26.80 ± 17.36*** | 37.50 ± 17.99*** | 23.25 ± 14.57** |
| 52 | Quinolone | Cinoxacin | 15.33 ± 8.08*** | 13.00 ± 10.44** | 46.00 ± 52.99* |
| 53 | | Nalidixic acid | 43.00 ± 25.46*** | 2.00 ± 4.00 | 0.00 ± 0.00 |
| 54 | Rifamycin | Rifampicin | 67.50 ± 7.55*** | 159.00 ± 30.25*** | 191.00 ± 90.60*** |

| Sr. No. | Class | Antibiotic | *Pseudomonas aeruginosa* | | |
|---|---|---|---|---|---|
| | | | ***In vivo* efficacy** (worm activity count) (Mean ± SEM) | | |
| | | | **24 h end-point** | **48 h end-point** | **72 h end-point** |
| 55 | Spiropyrimidinetrione | Zoliflodacin | 50.00 ± 28.58*** | 0.75 ± 1.50 | 18.33 ± 3.21*** |
| 56 | Streptogramin | Virginiamycin complex | 8.60 ± 4.22** | 0.00 ± 0.00 | 5.25 ± 7.09 |
| 57 | Sulfonamide | Sulfonilamide | 30.50 ± 20.37*** | 12.00 ± 0.00** | 32.00 ± 9.85*** |
| 58 | Sulfone antimicrobial | Dapsone | 3.25 ± 4.27 | 3.00 ± 6.00 | 2.75 ± 3.40 |
| 59 | Synthetic biocide / FabI inhibitor | Triclosan | 0.40 ± 0.89 | 7.00 ± 9.45 | 10.25 ± 12.39 |
| 60 | Tetracycline | Doxycycline Hydrochloride | 24.60 ± 7.70*** | 92.00 ± 10.15*** | 126.00 ± 19.08*** |
| 61 | | Lymecycline | 21.33 ± 4.93*** | 0.00 ± 0.00 | 0.00 ± 0.00 |
| 62 | | Meclocycline | 27.00 ± 14.42*** | 85.00 ± 19.80*** | 13.00 ± 4.36** |
| 63 | | Oxytetracycline | 30.00 ± 7.00*** | 38.50 ± 19.09*** | 17.00 ± 10.15** |
| 64 | | Rolitetracycline | 14.67 ± 5.69*** | 0.00 ± 0.00 | 0.00 ± 0.00 |
| 65 | Thiopeptide antibiotic | Nosiheptide | 39.00 ± 16.91*** | 77.00 ± 10.58*** | 93.00 ± 30.64*** |
| 66 | Cephalosporin | Cefadroxil | 15.33 ± 4.73*** | 16.00 ± 2.00*** | 30.00 ± 13.00*** |
| 67 | | Cefaclor | 7.60 ± 6.99 | 2.67 ± 4.62 | 6.00 ± 6.56 |
| 68 | | Cefixime | 21.00 ± 2.00*** | 18.33 ± 9.61*** | 95.00 ± 23.64*** |
| 69 | | Cefotaxime | 18.75 ± 6.70*** | 73.67 ± 27.93*** | 127.67 ± 28.75*** |
| 70 | | Ceftazidine | 3.75 ± 4.35 | 0.00 ± 0.00 | 1.67 ± 2.89 |
| 71 | | Ceftriaxone | 21.50 ± 2.12*** | 29.50 ± 6.36*** | 83.33 ± 36.25*** |
| 72 | β-lactam / aminopenicillin | Amoxicillin | 35.33 ± 15.28*** | 46.00 ± 21.21*** | 134.00 ± 5.66*** |
| 73 | | Ampicillin | 44.33 ± 10.69*** | 60.67 ± 10.69*** | 151.67 ± 15.18*** |
| 74 | β-lactam / carbapenem | Meropenem | 8.25 ± 6.65* | 4.40 ± 4.56 | 0.00 ± 0.00 |
| 75 | | Imipenem | 69.00 ± 35.36*** | 37.75 ± 12.12*** | 60.00 ± 42.95** |

None of the antibiotics, except triclosan, showed toxicity towards worms at the tested concentration till 72 h.

**Table S2 (c). Results of *in vivo* anti-pathogenic assay with all the 74 antibiotics against *Clostridioides difficile***

| Sr. No. | Class | Antibiotic | ***Clostridioides difficile*** | | |
|---|---|---|---|---|---|
| | | | ***In vivo* efficacy** (worm activity count) (Mean ± SEM) | | |
| | | | **24 h end-point** | **48 h end-point** | **72 h end-point** |
| 1 | Aminoglycoside | Amikacin | 0.00 ± 0.00*** | 0.00 ± 0.00 | 0.33 ± 0.82 |
| 2 | | Kanamycin | 0.00 ± 0.00 | 0.00 ± 0.00 | 0.00 ± 0.00 |
| 3 | | Streptomycin | 0.00 ± 0.00*** | 0.00 ± 0.00 | 0.00 ± 0.00 |
| 4 | | Tobramycin | 48.00 ± 16.97 | 58.50 ± 82.73 | 0.00 ± 0.00 |
| 5 | Aminomethylcycline / tetracycline | Omadacycline | 0.00 ± 0.00 | 0.00 ± 0.00 | 0.00 ± 0.00 |
| 6 | Amphenicol | Chloramphenicol | 25.80 ± 12.74* | 34.60 ± 47.39 | 0.00 ± 0.00 |
| 7 | | Florfenicol | 34.33 ± 17.77* | 20.20 ± 30.30 | 47.83 ± 64.35 |
| 8 | | thiamphenicol | 14.50 ± 14.86 | 35.00 ± 48.73 | 0.00 ± 0.00 |
| 9 | Antifolate / diaminopyrimidine | Trimethoprim | 4.50 ± 6.25 | 2.00 ± 4.47 | 0.67 ± 1.63 |
| 10 | | Iclaprim | 17.33 ± 8.43* | 56.80 ± 76.05 | 36.33 ± 72.40 |
| 11 | Antimycobacterial – ethambutol class | Ethambutol | 0.67 ± 1.63** | 0.80 ± 1.79 | 9.83 ± 21.73 |
| 12 | Antimycobacterial – hydrazide | Isoniazid | 2.17 ± 5.31 | 0.00 ± 0.00 | 1.33 ± 3.27 |
| 13 | Antimycobacterial – pyrazinamide class | Pyrazinamide | 0.00 ± 0.00 | 0.00 ± 0.00 | 0.00 ± 0.00 |
| 14 | Bisphenol / antiseptic; FabI inhibitor | Triclosan | 23.17 ± 20.03 | 0.00 ± 0.00 | 0.67 ± 1.63 |
| 15 | Boron-containing antimicrobial / leucyl-tRNA synthetase inhibitor | Epetraborole | 41.33 ± 12.42* | 48.00 ± 19.80 | 0.00 ± 0.00 |
| 16 | Coumarin antibiotic | Novobiocin | 5.50 ± 0.71 | 0.00 ± 0.00 | 0.00 ± 0.00 |
| 17 | Cyclic lipopeptide / biosurfactant | Surfactin | 1.50 ± 3.67 | 1.80 ± 4.02 | 0.83 ± 2.04 |
| 18 | Diarylpyridine | Bedaquiline | 0.00 ± 0.00*** | 1.00 ± 2.24 | 1.33 ± 3.27 |
| 19 | Experimental antimicrobial compound | U105920 | 0.00 ± 0.00*** | 1.00 ± 2.24 | 11.17 ± 22.81 |
| 20 | | AZ-1 | 1.67 ± 4.08 | 0.00 ± 0.00 | 0.00 ± 0.00 |
| 21 | Experimental LpxC inhibitor | CHIR-090 | 8.33 ± 11.47 | 0.00 ± 0.00 | 0.67 ± 1.63 |
| 22 | | PF-5081090 | 8.67 ± 9.67 | 0.00 ± 0.00 | 0.00 ± 0.00 |
| 23 | Fluoroquinolone | Delafloxacin | 35.33 ± 18.27* | 44.80 ± 33.30 | 86.17 ± 69.33 |

| Sr. No. | Class | Antibiotic | ***Clostridioides difficile*** | | |
|---|---|---|---|---|---|
| | | | ***In vivo* efficacy** (worm activity count) (Mean ± SEM) | | |
| | | | **24 h end-point** | **48 h end-point** | **72 h end-point** |
| 24 | | Gatifloxacin | 30.33 ± 21.21 | 50.75 ± 24.66** | 56.33 ± 77.00 |
| 25 | | Levofloxacin | 31.20 ± 12.52** | 0.50 ± 1.00 | 2.17 ± 5.31 |
| 26 | | Lomefloxacin | 32.17 ± 19.92* | 25.40 ± 40.92 | 26.33 ± 64.50 |
| 27 | | Moxifloxacin | 32.20 ± 8.07*** | 37.00 ± 22.32* | 28.00 ± 26.98 |
| 28 | | Norfloxacin | 25.00 ± 11.05** | 48.00 ± 68.06 | 64.00 ± 100.40 |
| 29 | | Ofloxacin | 37.83 ± 14.47** | 16.50 ± 12.02 | 60.17 ± 89.45 |
| 30 | | Sparfloxacin | 43.50 ± 8.57*** | 18.33 ± 8.02** | 70.80 ± 41.22* |
| 31 | | Ciprofloxacin | 38.20 ± 17.98** | 78.00 ± 49.67* | 68.00 ± 104.97 |
| 32 | Glycopeptide | Teicoplanin complex | 13.66 ± 15.62 | 0.00 ± 0.00 | 0.67 ± 1.63 |
| 33 | Glycopeptide/lipoglycopeptide-related | Ramoplanin complex | 17.83 ± 17.58 | 0.00 ± 0.00 | 6.17 ± 8.77 |
| 34 | Glycylcycline / tetracycline | Tigecycline | 0.00 ± 0.00 | 0.00 ± 0.00 | 0.00 ± 0.00 |
| 35 | Ketolide | Telithromycin | 12.00 ± 11.54 | 3.60 ± 3.78 | 12.67 ± 24.06 |
| 36 | Lipopeptide | Daptomycin | 0.00 ± 0.00 | 1.00 ± 1.41 | 0.00 ± 0.00 |
| 37 | Macrocyclic peptide | Murepavidin | 0.00 ± 0.00 | 0.00 ± 0.00 | 0.00 ± 0.00 |
| 38 | Macrolide | Clarithromycin | 31.00 ± 10.44** | 0.00 ± 0.00 | 4.33 ± 8.80 |
| 39 | | Erythromycin | 28.50 ± 8.96** | 20.80 ± 28.91 | 7.00 ± 10.86 |
| 40 | Macrolide – azalide | Azithromycin | 43.00 ± 8.49* | 0.00 ± 0.00 | 12.67 ± 19.03 |
| 41 | Nitroimidazole antimycobacterial | Pretomanid | 2.00 ± 4.90 | 0.00 ± 0.00 | 4.17 ± 10.21 |
| 42 | Oxazolidinone | Linezolid | 15.33 ± 17.61 | 0.00 ± 0.00 | 15.60 ± 24.17 |
| 43 | | Radezolid | 23.33 ± 24.43 | 0.00 ± 0.00 | 18.33 ± 28.65 |
| 44 | | Sutezolid | 33.00 ± 7.07* | 0.00 ± 0.00 | 0.00 ± 0.00 |
| 45 | | Tedizolid | 19.50 ± 28.72 | 0.00 ± 0.00 | 0.00 ± 0.00 |
| 46 | Pleuromutilin | Azamulin | 15.17 ± 16.86 | 8.80 ± 19.68 | 23.33 ± 36.29 |
| 47 | | Lefamulin | 12.33 ± 13.17 | 0.00 ± 0.00 | 0.00 ± 0.00 |
| 48 | | Retapamulin | 25.00 ± 23.93 | 0.00 ± 0.00 | 0.67 ± 1.63 |
| 49 | | Tiamulin | 9.00 ± 9.03 | 0.00 ± 0.00 | 0.00 ± 0.00 |
| 50 | | Valnemulin | 17.50 ± 22.01 | 2.00 ± 2.83 | 2.17 ± 3.71 |
| 51 | Polymyxin / cyclic lipopeptide | Polymyxin B | 16.67 ± 13.43 | 14.40 ± 17.64 | 10.83 ± 13.95 |
| 52 | Quinolone | Cinoxacin | 15.17 ± 13.26 | 40.60 ± 58.67 | 65.83 ± 111.06 |
| 53 | | Nalidixic acid | 0.00 ± 0.00*** | 0.00 ± 0.00 | 0.00 ± 0.00 |
| 54 | Rifamycin | Rifampicin | 43.00 ± 15.86** | 29.80 ± 35.24 | 0.50 ± 1.22 |

| Sr. No. | Class | Antibiotic | ***Clostridioides difficile*** | | |
|---|---|---|---|---|---|
| | | | ***In vivo* efficacy** (worm activity count) (Mean ± SEM) | | |
| | | | **24 h end-point** | **48 h end-point** | **72 h end-point** |
| 55 | Spiropyrimidinetrione | Zoliflodacin | 17.67 ± 12.55 | 1.20 ± 2.68 | 11.33 ± 23.63 |
| 56 | Streptogramin | Virginiamycin complex | 34.00 ± 7.94** | 0.00 ± 0.00 | 0.00 ± 0.00 |
| 57 | Sulfonamide | Sulfonilamide | 0.67 ± 1.63** | 2.20 ± 3.49 | 0.67 ± 1.63 |
| 58 | Sulfonamide / sulfones | Dapsone | 0.67 ± 1.63** | 0.00 ± 0.00 | 2.33 ± 5.72 |
| 59 | Tetracycline | Doxycycline Hydrochloride | 40.00 ± 24.38* | 39.40 ± 19.46** | 50.20 ± 54.11 |
| 60 | | Lymecycline | 0.00 ± 0.00 | 2.00 ± 2.83 | 0.00 ± 0.00 |
| 61 | | Meclocycline | 0.00 ± 0.00 | 0.00 ± 0.00 | 0.00 ± 0.00 |
| 62 | | Oxytetracycline | 16.00 ± 13.37 | 0.00 ± 0.00 | 0.00 ± 0.00 |
| 63 | | Rolitetracycline | 0.00 ± 0.00 | 0.00 ± 0.00 | 0.00 ± 0.00 |
| 64 | Thiopeptide antibiotic | Nosiheptide | 8.00 ± 7.62 | 4.80 ± 8.23 | 16.50 ± 23.71 |
| 65 | β-lactam – aminopenicillin | Amoxicillin | 15.83 ± 27.14 | 0.80 ± 1.79 | 9.00 ± 17.23 |
| 66 | | Ampicillin | 0.00 ± 0.00 | 0.00 ± 0.00 | 3.83 ± 6.15 |
| 67 | β-lactam – carbapenem | Meropenem | 0.00 ± 0.00 | 0.00 ± 0.00 | 0.00 ± 0.00 |
| 68 | | Imipenem | 0.00 ± 0.00 | 0.00 ± 0.00 | 0.00 ± 0.00 |
| 69 | Cephalosporin | Cefadroxil | 0.00 ± 0.00 | 0.00 ± 0.00 | 0.00 ± 0.00 |
| 70 | | Cefaclor | 10.50 ± 9.50 | 0.80 ± 1.79 | 0.00 ± 0.00 |
| 71 | | Cefixime | 0.00 ± 0.00 | 1.50 ± 2.12 | 0.00 ± 0.00 |
| 72 | | Cefotaxime | 22.00 ± 3.61*** | 0.00 ± 0.00 | 2.00 ± 4.90 |
| 73 | | Ceftazidine | 15.40 ± 4.56** | 4.80 ± 8.67 | 2.33 ± 5.72 |
| 74 | | Ceftriaxone | 34.67 ± 12.50** | 0.00 ± 0.00 | 0.00 ± 0.00 |

None of the antibiotics, except triclosan, showed toxicity towards worms at the tested concentration till 72 h.

### Table S3. Antibiotic efficacy against MRSA

| | Group | Treatment | Worm Activity (Mean ± SEM) | |
|---|---|---|---|---|
| | | | **24 h endpoint** | **48 h endpoint** |
| | A | Health control | 28.83 ± 6.88 | 28.16 ± 6.61 $^{a}$ |
| | B | Infection control | 4.83 ± 4.30 $^{adefg}$ | 0 ± 0 $^{a}$ |
| | C | Vehicle control | 4.5 ± 3.78 $^{adefg}$ | 0 ± 0 $^{a}$ |
| Aminoglycosides | D | Streptomycin | 22.67 ± 6.11 $^{bc}$ | 0.00 ± 0.00 $^{a}$ |
| | E | Tobramycin | 20.67 ± 10.95 $^{bc}$ | 0.00 ± 0.00 $^{a}$ |
| | F | Amikacin | 19.83 ± 8.50 $^{bc}$ | 0.00 ± 0.00 $^{a}$ |
| | G | Kanamycin | 21.17 ± 8.82 $^{bc}$ | 0.00 ± 0.00 $^{a}$ |
| | A | Health control | 28.83 ± 6.88 $^{bc}$ | 28.16 ± 6.61 $^{gh}$ |
| | B | Infection control | 4.83 ± 4.30 $^{adefgij}$ | 0 ± 0 $^{adi}$ |
| | C | Vehicle control | 4.5 ± 3.78 $^{adefgij}$ | 0 ± 0 $^{adi}$ |
| Fluoroquinolones | D | Sparfloxacin | 34.67 ± 15.44 $^{bch}$ | 34.50 ± 16.37 $^{bcgh}$ |
| | E | Gatifloxacin | 34.00 ± 13.17 $^{bch}$ | 31.40 ± 19.15 $^{bcgh}$ |
| | F | Ofloxacin | 38.00 ± 17.41 $^{bch}$ | 16.17 ± 12.97 |
| | G | Norfloxacin | 31.83 ± 16.96 $^{bch}$ | 3.67 ± 4.23 $^{adej}$ |
| | H | Ciprofloxacin | 7.20 ± 4.15 $^{defgij}$ | 3.50 ± 5.43 $^{adej}$ |
| | I | Levofloxacin | 34.25 ± 3.30 $^{bch}$ | 23.40 ± 8.88 $^{bc}$ |
| | J | Moxifloxacin | 40.50 ± 18.02 $^{bch}$ | 31.00 ± 11.58 $^{bcgh}$ |
| | A | Health control | 28.83 ± 6.88 $^{bc}$ | 28.16 ± 6.61 $^{bcdef}$ |
| | B | Infection control | 4.83 ± 4.30 $^{adef}$ | 0 ± 0 $^{a}$ |
| | C | Vehicle control | 4.5 ± 3.78 $^{adef}$ | 0 ± 0 $^{a}$ |
| Cephalosporin | D | Ceftriaxone | 22.83 ± 14.92 $^{bc}$ | 4.33 ± 4.04 $^{a}$ |
| | E | Cephotaxime | 17.75 ± 5.74 $^{bc}$ | 1.00 ± 2.00 $^{a}$ |
| | F | Cefixime | 20.50 ± 4.68 $^{bc}$ | 6.00 ± 5.83 $^{a}$ |
| | A | Health control | 28.83 ± 6.88 $^{bc}$ | 28.16 ± 6.61 $^{bcdef}$ |
| | B | Infection control | 4.83 ± 4.30 $^{adef}$ | 0 ± 0 $^{a}$ |
| | C | Vehicle control | 4.5 ± 3.78 $^{adef}$ | 0 ± 0 $^{a}$ |
| Macrolide | D | Erythromycin | 21.00 ± 13.08 $^{bc}$ | 2.00 ± 4.90 $^{a}$ |
| | E | Azithromycin | 22.50 ± 7.42 $^{bc}$ | 0.00 ± 0.00 $^{a}$ |
| | F | Clarithromycin | 28.00 ± 11.96 $^{bc}$ | 0.00 ± 0.00 $^{a}$ |
| | A | Health control | 28.83 ± 6.88 $^{bcde}$ | 28.16 ± 6.61 $^{defghk}$ |
| | B | Infection control | 4.83 ± 4.30 $^{afghijl}$ | 0 ± 0 $^{i,l}$ |
| | C | Vehicle control | 4.5 ± 3.78 $^{afghijl}$ | 0 ± 0 $^{i,l}$ |
| Rest classes | D | Nalidixic acid | 6.80 ± 9.34 $^{afghijl}$ | 3.00 ± 7.35 $^{i,l}$ |
| | E | Imipenem | 7.80 ± 8.50 $^{afghijl}$ | 0.00 ± 0.00 $^{i,l}$ |
| | F | Ampicillin | 38.33 ± 21.20 $^{bcde}$ | 0.67 ± 1.63 $^{i,l}$ |
| | G | Teicoplanin | 31.40 ± 11.87 $^{bcdej}$ | 0.67 ± 1.63 $^{i,l}$ |
| | H | Linezolid | 37.25 ± 3.95 $^{bcde}$ | 0.00 ± 0.00 $^{i,l}$ |
| | I | Doxycycline Hydrochloride | 36.50 ± 11.29 $^{bcdej}$ | 35.50 ± 17.72 $^{bcdefghk}$ |
| | J | Chloramphenicol | 56.00 ± 16.24 $^{bcdegikl}$ | 28.25 ± 8.42 |
| | K | Novobiocin | 23.40 ± 10.95 $^{j}$ | 0.00 ± 0.00 $^{i,l}$ |
| | L | Rifampicin | 30.50 ± 11.47 $^{bcdej}$ | 27.80 ± 12.97 |

Superscript letters denote statistically significant differences ($P < 0.05$) from the indicated group within the same column based on Tukey's multiple comparisons test following two-way ANOVA. a indicates significantly different from Group A; b from Group B; c from Group C; d from Group D; e from Group E; f from Group F; g from Group G. Seventy-two-hour endpoint is not included here, as some of the antibiotic wells registered higher worm activity owing to progeny worms at this time point.

### Table S4. Antibiotic efficacy against *P. aeruginosa*

| | Group | Treatment | **Worm Activity** (Mean ± SEM) | |
|---|---|---|---|---|
| | | | **24 h endpoint** | **48 h endpoint** |
| | A | Health control | 39.1 ± 15.84 $^{bcdefg}$ | 49.3 ± 9.66 $^{bcdfg}$ |
| | B | Infection control | 3.2 ± 4.21$^{a}$ | 1.6 ± 2.06$^{e}$ |
| Aminoglycosides | C | Vehicle control | 2.4 ± 3.09$^{a}$ | 2.7 ± 3.71$^{e}$ |
| | D | Streptomycin | 6.00 ± 5.20$^{a}$ | 1.50 ± 1.91$^{e}$ |
| | E | Tobramycin | 1.33 ± 2.31$^{a}$ | 50.33 ± 10.07 $^{bcdfg}$ |
| | F | Amikacin | 2.00 ± 3.46$^{a}$ | 3.20 ± 4.60$^{e}$ |
| | G | Kanamycin | 0.00 ± 0.00$^{a}$ | 2.80 ± 4.38$^{e}$ |
| | A | Health control | 39.10 ± 15.84$^{bei}$ | 49.30 ± 9.66$^{bcdf}$ |
| | B | Infection control | 3.20 ± 4.21$^{af}$ | 1.60 ± 2.06$^{adehj}$ |
| Fluoroquinolones | C | Vehicle control | 2.40 ± 3.09$^{af}$ | 2.70 ± 3.71$^{adehj}$ |
| | D | Sparfloxacin | 27.25 ± 14.82$^{f}$ | 63.80 ± 31.06$^{bcgh}$ |
| | E | Gatifloxacin | 13.00 ± 1.73$^{a}$ | 56.25 ± 13.87$^{bcgh}$ |
| | F | Ofloxacin | 28.00 ± 11.97$^{bc}$ | 70.50 ± 15.95$^{ghi}$ |
| | G | Norfloxacin | 23.00 ± 17.44 | 0.00 ± 0.00$^{befi}$ |
| | H | Ciprofloxacin | 20.00 ± 7.07 | 26.00 ± 16.52$^{dgj}$ |
| | I | Levofloxacin | 20.25 ± 16.80 | 30.80 ± 15.01$^{bcg}$ |
| | J | Moxifloxacin | 9.50 ± 5.20$^{a}$ | 62.67 ± 20.84$^{cdgj}$ |
| | A | Health control | 39.10 ± 15.84$^{bcde}$ | 49.30 ± 9.66$^{bcdf}$ |
| | B | Infection control | 3.20 ± 4.21$^{aef}$ | 1.60 ± 2.06$^{adef}$ |
| Cephalosporin | C | Vehicle control | 2.40 ± 3.09$^{adef}$ | 2.70 ± 3.71$^{adef}$ |
| | D | Ceftriaxone | 21.50 ± 2.12$^{ac}$ | 29.50 ± 6.36$^{abc}$ |
| | E | Cefotaxime | 18.75 ± 6.70$^{abc}$ | 73.67 ± 27.93$^{bcdf}$ |
| | F | Cefixime | 21.00 ± 2.00$^{abc}$ | 18.33 ± 9.61$^{abce}$ |
| | A | Health control | 39.10 ± 15.84$^{bc}$ | 49.30 ± 9.66$^{bcd}$ |
| | B | Infection control | 3.20 ± 4.21$^{def}$ | 1.60 ± 2.06$^{ef}$ |
| Macrolide | C | Vehicle control | 2.40 ± 3.09$^{def}$ | 2.70 ± 3.71$^{ef}$ |
| | D | Erythromycin | 25.50 ± 12.66$^{bc}$ | 0.00 ± 0.00$^{aef}$ |
| | E | Azithromycin | 48.50 ± 3.54$^{bc}$ | 45.00 ± 20.66 |
| | F | Clarithromycin | 46.67 ± 3.06$^{bc}$ | 50.75 ± 18.73$^{bcd}$ |
| | A | Health control | 39.10 ± 15.84$^{bcdfhj}$ | 49.30 ± 9.66$^{bcdfhj}$ |
| | B | Infection control | 3.20 ± 4.21$^{adefhj}$ | 1.60 ± 2.06$^{adefhj}$ |
| Rest classes | C | Vehicle control | 2.40 ± 3.09$^{adefhj}$ | 2.70 ± 3.71$^{adefhj}$ |
| | D | Nalidixic acid | 43.00 ± 25.46$^{bcfh}$ | 2.00 ± 4.00$^{abcfhj}$ |
| | E | Imipenem | 69.00 ± 35.36$^{abdfghj}$ | 37.75 ± 12.12$^{j}$ |
| | F | Ampicillin | 44.33 ± 10.69$^{bcdh}$ | 60.67 ± 10.69$^{hj}$ |
| | G | Teicoplanin | 2.75 ± 2.22$^{aefhj}$ | 4.00 ± 5.66$^{abcdfhj}$ |
| | H | Linezolid | 28.50 ± 6.36$^{aj}$ | 43.50 ± 3.54$^{befj}$ |
| | I | Doxycycline (hydrochloride) | 24.60 ± 7.70$^{bcj}$ | 92.00 ± 10.15$^{abcefghj}$ |
| | J | Chloramphenicol | 53.00 ± 15.64$^{bcg}$ | 4.00 ± 6.73$^{abcdfj}$ |
| | K | Novobiocin | 6.50 ± 5.97$^{adfj}$ | 33.00 ± 6.56$^{b}$ |
| | L | Rifampicin | 67.50 ± 7.55$^{abcghik}$ | 159.00 ± 30.25$^{abcdefghijk}$ |

Meaning of superscripts: Groups sharing at least one superscript letter are not significantly different (Tukey's multiple-comparison test, P > 0.05). Groups with no superscript letter in common differ significantly (P < 0.05). Seventy-two-hour endpoint is not included here, as some of the antibiotic wells registered higher worm activity owing to progeny worms at this time point.

### Table S5. Antibiotic efficacy against *Cl. difficile*

| | Group | Treatment | **Worm Activity** (Mean ± SEM) | |
|---|---|---|---|---|
| | | | **24 h endpoint** | **48 h endpoint** |
| | A | Health control | 29.36 ± 9.73$^{a}$ | 21.77 ± 12.88$^{a}$ |
| | B | Infection control | 2.94 ± 1.18$^{b}$ | 0.00 ± 0.00$^{b}$ |
| | C | Vehicle control | 2.88 ± 1.95$^{b}$ | 0.00 ± 0.00$^{b}$ |
| Aminoglycosides$^{\#}$ | D | Streptomycin | 0.00 ± 0.00$^{b}$ | 0.00 ± 0.00$^{b}$ |
| | E | Tobramycin | 48.00 ± 16.97$^{a}$ | 58.50 ± 82.73$^{a}$ |
| | F | Amikacin | 0.00 ± 0.00$^{b}$ | 0.00 ± 0.00$^{b}$ |
| | G | Kanamycin | 0.00 ± 0.00$^{b}$ | 0.00 ± 0.00$^{b}$ |
| | A | Health control | 29.36 ± 9.73$^{ab}$ | 21.77 ± 12.88$^{ab}$ |
| | B | Infection control | 2.94 ± 1.18$^{c}$ | 0.00 ± 0.00$^{a}$ |
| | C | Vehicle control | 2.88 ± 1.95$^{bc}$ | 0.00 ± 0.00$^{a}$ |
| Fluoroquinolones | D | Sparfloxacin | 43.50 ± 8.57$^{a}$ | 18.33 ± 8.02$^{ab}$ |
| | E | Gatifloxacin | 30.33 ± 21.21$^{ab}$ | 50.75 ± 24.66$^{bc}$ |
| | F | Ofloxacin | 37.83 ± 14.46$^{a}$ | 16.50 ± 12.02$^{ab}$ |
| | G | Norfloxacin | 25.00 ± 11.04$^{abc}$ | 0.00 ± 0.00$^{a}$ |
| | H | Ciprofloxacin | 38.20 ± 17.97$^{a}$ | 78.00 ± 49.66$^{c}$ |
| | I | Levofloxacin | 31.20 ± 12.51$^{ab}$ | 0.50 ± 1.00$^{a}$ |
| | J | Moxifloxacin | 32.20 ± 8.07$^{ab}$ | 22.31$^{abc}$ |
| | A | Health control | 29.36 ± 9.73$^{a}$ | 21.77 ± 12.88$^{a}$ |
| | B | Infection control | 2.94 ± 1.18$^{c}$ | 0.00 ± 0.00$^{b}$ |
| | C | Vehicle control | 2.88 ± 1.95$^{c}$ | 0.00 ± 0.00$^{b}$ |
| Cephalosporin | D | Ceftriaxone | 34.67 ± 12.50$^{a}$ | 0.00 ± 0.00$^{b}$ |
| | E | Cephotaxime | 22.00 ± 3.61$^{b}$ | 0.00 ± 0.00$^{b}$ |
| | F | Cefixime | 0.00 ± 0.00$^{c}$ | 1.50 ± 2.12$^{b}$ |
| | A | Health control | 29.36 ± 9.73$^{a}$ | 21.77 ± 12.88$^{a}$ |
| | B | Infection control | 2.94 ± 1.18$^{b}$ | 0.00 ± 0.00$^{b}$ |
| | C | Vehicle control | 2.88 ± 1.95$^{b}$ | 0.00 ± 0.00$^{b}$ |
| Macrolide | D | Erythromycin | 28.50 ± 8.96$^{a}$ | 0.00 ± 0.00$^{b}$ |
| | E | Azithromycin | 43.00 ± 8.49$^{a}$ | 0.00 ± 0.00$^{b}$ |
| | F | Clarithromycin | 31.00 ± 10.44$^{a}$ | 0.00 ± 0.00$^{b}$ |
| | A | Health control | 29.36 ± 9.73$^{ab}$ | 21.77 ± 12.88$^{a}$ |
| | B | Infection control | 2.94 ± 1.18$^{c}$ | 0 ± 0$^{b}$ |
| | C | Vehicle control | 2.88 ± 1.95$^{c}$ | 0 ± 0$^{b}$ |
| Rest classes | D | Nalidixic acid | 0.00 ± 0.00$^{c}$ | 0.00 ± 0.00$^{ab}$ |
| | E | Imipenem | 0.00 ± 0.00$^{c}$ | 0.00 ± 0.00$^{ab}$ |
| | F | Ampicillin | 0.00 ± 0.00$^{bc}$ | 0.00 ± 0.00$^{ab}$ |
| | G | Teicoplanin | 13.66 ± 15.62$^{abc}$ | 0.00 ± 0.00$^{ab}$ |
| | H | Linezolid | 15.33 ± 17.61$^{abc}$ | 0.00 ± 0.00$^{ab}$ |
| | I | Doxycycline (hydrochloride) | 40.00 ± 24.38$^{a}$ | 39.40 ± 19.46$^{a}$ |
| | J | Chloramphenicol | 25.80 ± 12.73$^{abc}$ | 0.00 ± 0.00$^{b}$ |
| | K | Novobiocin | 5.50 ± 0.71$^{bc}$ | 0.00 ± 0.00$^{ab}$ |
| | L | Rifampicin | 43.00 ± 15.85$^{a}$ | 0.00 ± 0.00$^{b}$ |

Different superscript letters within the same column indicate statistically significant differences among treatments according to Tukey's multiple comparisons test following two-way ANOVA ($P < 0.05$), whereas values sharing at least one common superscript letter are not significantly different. Treatments are assigned Group A–L for identification only and do not represent statistical groupings. Seventy-two-hour endpoint is not included here, as some of the antibiotic wells registered higher worm activity owing to progeny worms at this time point. # = The only group that differs significantly from the others is Tobramycin at both 24 h and 48 h. All remaining groups are statistically similar to one another. Therefore, only two letters (a and b) are needed.

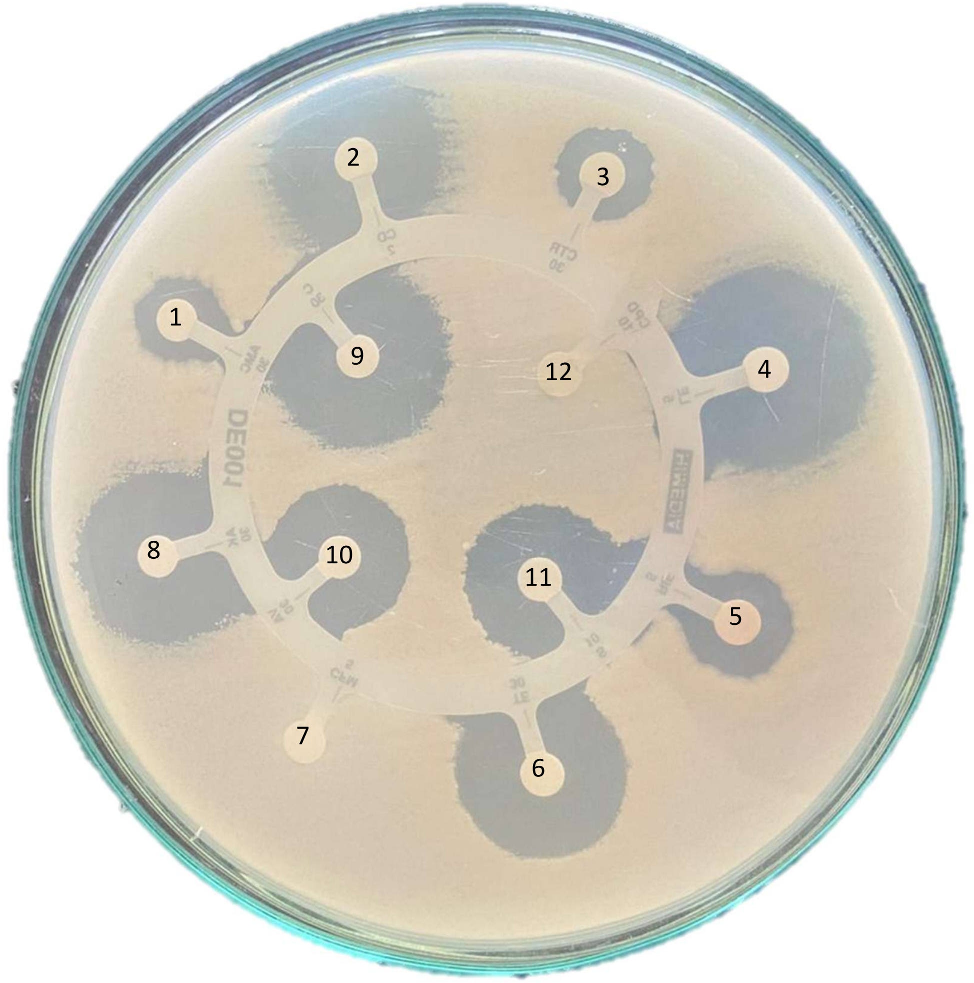

**Figure S1. Antibiotic Susceptibility profile of *MRSA*.** 1. Amikacin, 2. Amoxyclav, 3. Clindamycin, 4. Ceftriaxone, 5. Levofloxacin, 6. Rifampicin, 7. Tetracycline, 8. Cefixime, 9. Vancomycin, 10. Streptomycin, 11. Cefpodoxime, 12. Chloramphenicol.

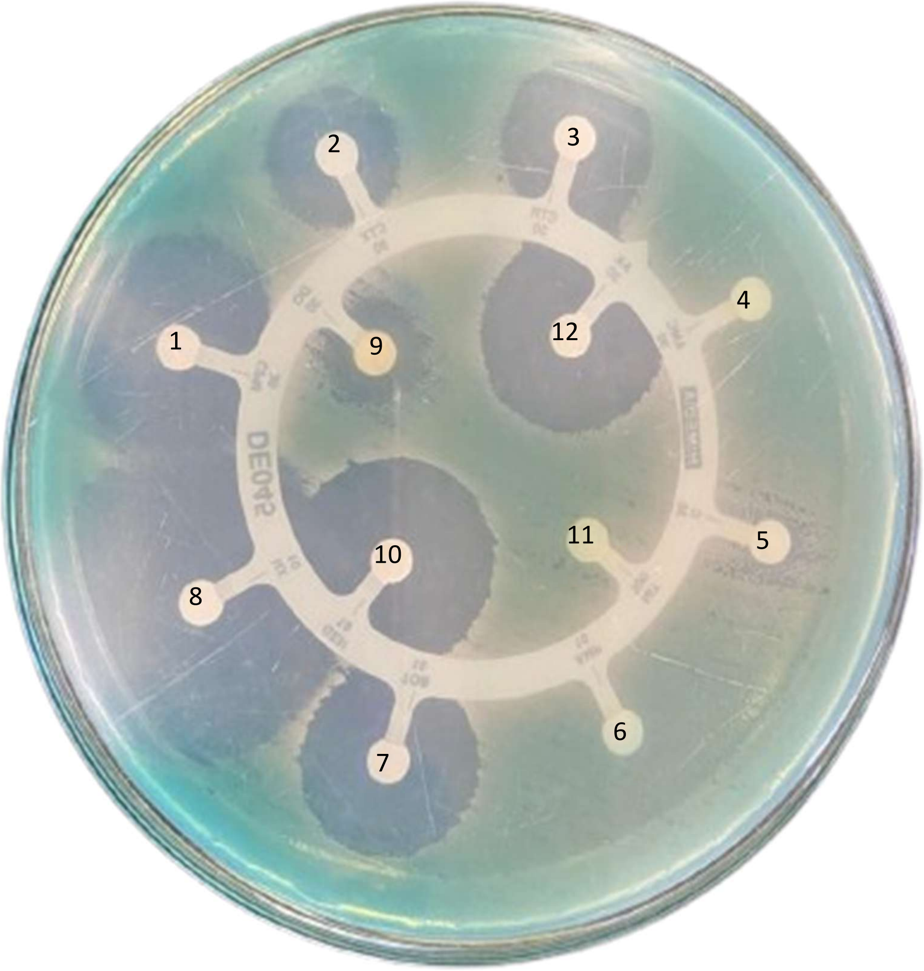

**Figure S2. Antibiotic Susceptibility profile of *P. aeruginosa*.** 1. Cefepime, 2. Cefotaxime, 3. Ceftriaxone, 4. Amoxyclav, 5. Chloramphenicol, 6. Ampicillin, 7. Tobramycin, 8. Norfloxacin, 9. Doxycycline Hydrochloride, 10. Gentamicin, 11. Nitrofurantoin, 12. Amikacin

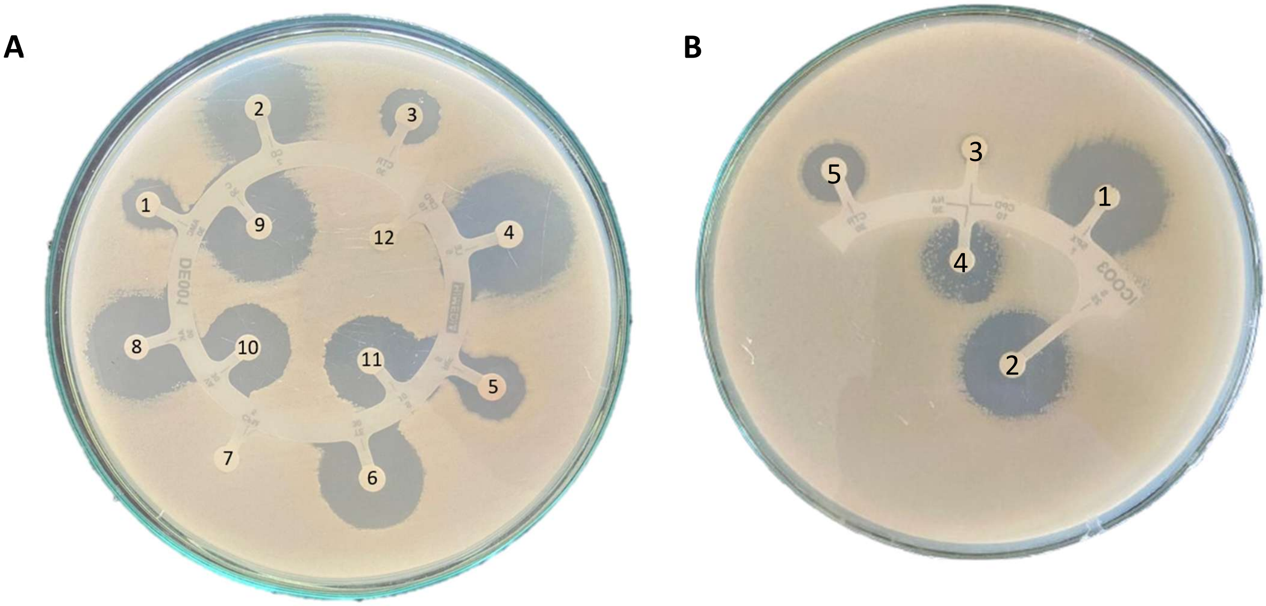

**Figure S3. Antibiotic Susceptibility profile of *Cl. difficile.* (A)** 1. Amoxyclav, 2. Clindamycin, 3. Ceftriaxone, 4. Levofloxacin, 5. Rifampicin, 6. Tetracycline, 7. Cefixime, 8. Amikacin, 9. Chloramphenicol, 10. Vancomycin, 11. Streptomycin, 12. Cefpodoxime. (**B)** 1. Sparfloxacin, 2. Streptomycin, 3. Cefpodoxime, 4. Nalidixic acid, 5. Ceftriaxone

### Appendix A
### Legend to Supplementary Videos (https://zenodo.org/records/24302220)

Videos were captured using Magnus Camera (5.1 MP) attached to LABOMED (Vision 2000) binocular microscope (halogen light source; 4X objective).

Video A-B: Health Control (HC): Worms in M9 buffer exposed neither to antibiotic nor bacterial pathogen, captured at 24 h (Video A) and 72 h (Video B)

**Methicillin-resistant *Staphylococcus aureus* (MRSA)**

Video C: Vehicle Control: Worms challenged with MRSA in presence of DMSO (0.15% v/v). DMSO did not interfere with bacterial virulence, and all worms were killed within 24 h.

Video D: Azithromycin though produced no zone of inhibition in disc diffusion assay, could confer significant protection on worms in face of MRSA challenge (24 h)

Video E: Erythromycin remained ineffective against MRSA during *in vitro* as well as *in vivo* assay. All worms challenged with MRSA in presence of erythromycin were killed within 24 hours.

Video F: Chloramphenicol showed anti-MRSA activity *in vitro* as well as *in vivo*. Worms challenged with MRSA in presence of chloramphenicol, not only survived, but also registered activity higher than the Health Control (24 h).

***Pseudomonas aeruginosa***

Video G: Vehicle Control: Worms challenged with *P. aeruginosa* in presence of DMSO (0.15% v/v). DMSO did not interfere with bacterial virulence, and all worms were killed within 24 h.

Video H1-H2: Though amikacin produced zone of inhibition during disc diffusion assay (DDA), its protective effect on pathogen-challenged worms was not visible till first 48 h (Video H1). Longer observation could identify partial protective effect of this antibiotic at 72 h time point (Video H2).

Video I1-I2: Cefotaxime was not only able to inhibit bacterial growth on agar plate as well as confer protection on worm population in face of pathogen challenge (Video I1; 48 h), but also allowed progeny production in pathogen-challenged worm population (Video I2; 72 h).

Video J: Chloramphenicol though produced a relatively smaller zone of inhibition on agar plate, could confer significant protection on worms in face of *P. aeruginosa* challenge (24 h).

Video K1-K2: Tobramycin was able to produce a zone of inhibition in DDA. However, when worms were challenged with *P. aeruginosa* in presence of tobramycin, this antibiotic conferred a delayed protection on worm population. Till first 24 hours, worms remained largely inactive under the pathogen-influence (Video K1), but 48 h onward antibiotic allowed not only worm recovery, but their reproduction too. Protective effect of tobramycin was sustained till 72 h (Video K2).

***Clostridioides difficile***

Video L: Vehicle Control: Worms challenged with *Cl. difficile* in presence of DMSO (0.15% v/v). DMSO did not interfere with bacterial virulence, and all worms were killed within 24 h.

Video M: Though amikacin proved effective in DDA, it could not protect worms in face of *Cl. difficile* challenge. All the worms challenged with *Cl. difficile* in presence of amikacin were killed within 24 hours.

Video N: *Cl. difficile* was found to be completely resistant to cefixime in DDA, and this matched with the ineffectiveness of this antibiotic *in vivo* too, when all the worms challenged with *Cl. difficile* in presence of cefixime were killed within 24 hours of pathogen-challenge.

Video O: Sparfloxacin produced a zone of inhibition in DDA, and was also able to confer a sustained protection on worm population challenged with *Cl. difficile* till 72 h.
